# A Molecular, Spatial, and Regulatory Atlas of the *Hydra vulgaris* Nervous System

**DOI:** 10.1101/2023.03.15.531610

**Authors:** Hannah Morris Little, Abby S. Primack, Jennifer Tsverov, Susanne Mühlbauer, Ben D. Cox, Christina Busse, Sandra Schneid, Amber Louwagie, Jack F. Cazet, Charles N. David, Jeffrey A. Farrell, Celina E. Juliano

**Affiliations:** Department of Molecular and Cellular Biology, University of California, Davis, CA 95616; Department of Biology, Ludwig-Maximilians-University Munich, 82152 Martinsried, Germany; Division of Developmental Biology, Eunice Kennedy Shriver National Institute of Child Health and Human Development, Bethesda, MD 20814, USA

## Abstract

*Hydra vulgaris*, a cnidarian with a simple nerve net, is an emerging model for developmental, regenerative, and functional neuroscience. Its genetic tractability and capacity for whole-system imaging make it well suited for studying neuron replacement, regeneration, and neural circuit function. Here, we present the most comprehensive molecular and spatial characterization of the *H. vulgaris* nervous system to date. Using single-cell RNA sequencing, we identified eight neuron types, each defined by distinct neuropeptide expression, and further resolved these into fifteen transcriptionally distinct subtypes with unique spatial distributions and morphologies. To investigate the gene regulatory networks underlying neuronal differentiation, we applied trajectory inference, identified key transcription factors, and performed ATAC-seq on sorted neurons to map chromatin accessibility. All datasets are available through an interactive, user-friendly web portal to support broad use by the research community. Together, these resources provide a foundation for uncovering molecular mechanisms that govern nervous system development, homeostasis, and regeneration in *H. vulgaris*.

**Summary Statement:** *Hydra vulgaris* is a model for regenerative and functional neuroscience. This study identifies fifteen neuron subtypes using scRNA-seq, maps spatial distributions, explores regulatory mechanisms, and provides an accessible web portal.

## Introduction

The small freshwater cnidarian *Hydra vulgaris* (**Fig. 1A**) is a classic model for studying regeneration due to its ability to regenerate its entire body, including the nervous system (Bode et al., 1988; Rentzsch et al., 2019; Trembley et al., 1744; Vogg et al., 2019b). Even in uninjured animals, neurogenesis pathways remain continuously active, enabling constant neuron replacement and providing easy experimental access to study the molecular mechanisms of neurogenesis (David and Gierer, 1974; David and Murphy, 1977; Siebert et al., 2019). Recently, *H. vulgaris* has also gained recognition as a valuable model in functional neuroscience, with the identification of neural circuits that drive specific behaviors (Dupre and Yuste, 2017; Giez et al., 2023; Yamamoto and Yuste, 2023). Building on this foundational research, *H. vulgaris* now serves as a robust model for regenerative neuroscience, offering tools to investigate the complete regeneration of an adult nervous system, from molecules to behavior.

**Figure 1.**
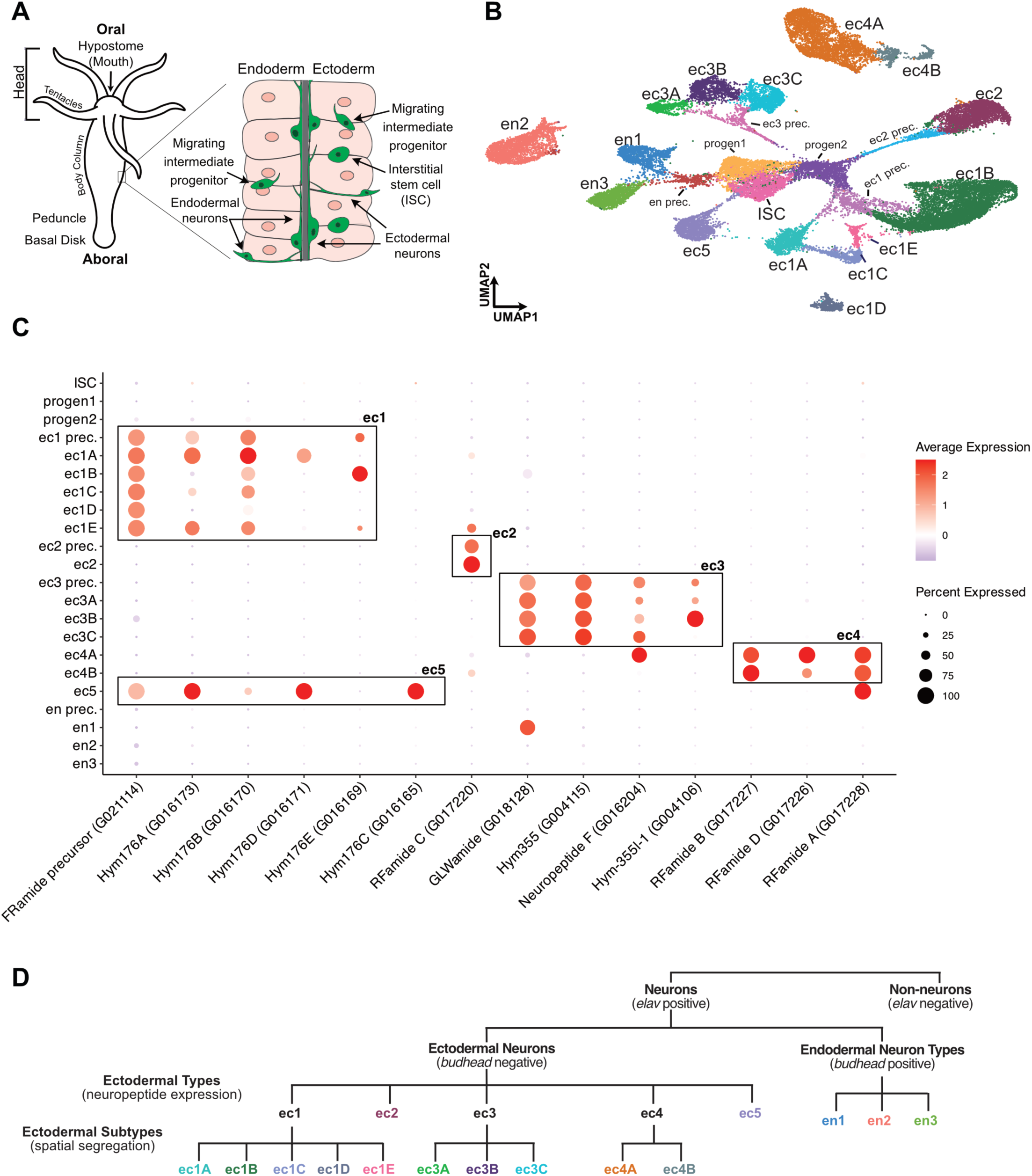
Single cell RNA sequencing reveals fifteen transcriptionally distinct neuron subtypes comprising the *Hydra vulgaris* nervous system. (A) The *Hydra* body is a radially symmetric, hollow tubular animal arranged around an oral-aboral axis. The oral end consists of the hypostome and tentacles (collectively referred to as the “head”), while the aboral end comprises the peduncle and basal disk. The body is formed by two epithelial monolayers, the endoderm and ectoderm, separated by an extracellular matrix. Neurons reside within the interstitial spaces between epithelial cells, forming two distinct nerve nets: one in the ectodermal layer and another in the endodermal layer (Keramidioti et al., 2024). Interstitial stem cells (ISCs), which give rise to neurons, are confined to the ectoderm, but intermediate neural progenitors migrate to generate endodermal neuron subtypes. **(B)** Single-cell RNA sequencing (scRNA-seq) was performed using two methods: Chromium Single-Cell Gene Expression (10x Genomics) (29,671 cells from this study) and Drop-Seq (5,400 cells from Siebert et al., 2019). A UMAP representation displays cell clusters annotated by their respective cell states: ISC (interstitial stem cell), progen (progenitor) prec (precursors), ec (ectodermal neurons), and en (endodermal neurons). **(C)** Dot plot showing the expression of neuropeptide genes across *Hydra vulgaris* neuron clusters. Dot color represents the average expression level of each gene within a cluster, while dot size indicates the percentage of cells in the cluster expressing the gene. The ec1, ec2, ec3, ec4, and ec5 neuron types have distinct neuropeptide expression patterns. **(D)** Schematic outlining the classification process for *H. vulgaris* neurons. First, neurons were identified in the scRNA-seq dataset by the expression of *elav*. Cells were further categorized into either the ectodermal or endodermal nerve net based on the expression of the endodermal marker *budhead*. Ectodermal neuron types were defined by distinct neuropeptide expression profiles, and ectodermal neuron subtypes were classified based on spatially restricted domains (see main text for detailed information on the classification scheme).

The *H. vulgaris* body is formed by two epithelial monolayers, an outer ectodermal layer and an inner endodermal layer (**Fig. 1A**). Its nervous system consists of approximately 3,000–5,000 neurons (around 3-5% of total body cells), organized into two distinct nerve nets, one within each epithelial layer (Bode et al., 1973; Keramidioti et al., 2024). *H. vulgaris* neurons are part of the interstitial cell lineage, and their replacement during both homeostasis and regeneration is supported by multipotent adult interstitial stem cells (ISCs) located among the ectodermal epithelial cells (**Fig. 1A**) (David and Gierer, 1974; David and Murphy, 1977). Due to passive tissue displacement toward the extremities, which leads to ongoing cell loss, ISCs continuously replenish neurons even in the absence of injury (Campbell, 1967; David and Gierer, 1974; Hager and David, 1997). The constant activity of neurogenesis pathways in uninjured adult polyps allows for the detailed profiling of differentiation states using single-cell RNA sequencing (scRNA-seq) (Siebert et al., 2019).

In addition to exhibiting continual, widespread neuronal regeneration, *H. vulgaris* are amenable to tests of gene function, such as gene knock-down through the expression of RNA hairpins or the electroporation of siRNAs (Juliano et al., 2014a; Lommel et al., 2017; Vogg et al., 2019a). Additionally, creating stable transgenic *H. vulgaris* lines is straightforward using cell-type-specific promoters (Siebert et al., 2008; Siebert et al., 2019; Wittlieb et al., 2006). This has enabled the expression of fluorescent calcium reporters like GCaMP to monitor neuronal activity and the use of nitroreductase to selectively ablate specific neurons (Dupre and Yuste, 2017; Giez et al., 2023; Noro et al., 2021). Together, these attributes make *H. vulgaris* an accessible model for investigating the molecular mechanisms underlying nervous system development and regeneration, from cellular differentiation to the formation of neural circuits driving behavior. Additionally, as a member of Cnidaria, the sister group to bilaterians, *H. vulgaris* provides a unique opportunity to explore the evolution of the molecular processes involved in nervous system development, regeneration, and function.

Decades of research have established a foundational framework for studying the *H. vulgaris* nervous system. Our basic understanding includes the source of new neurons, the relative distribution of neurons along the oral-aboral axis, and neurogenesis rates in an uninjured *H. vulgaris* (Bode et al., 1973; Bode et al., 1988; David, 2012; David and Gierer, 1974). In our previous work, where we created a whole-animal single-cell expression map, we transcriptionally profiled *H. vulgaris* neurons and largely analyzed in situ hybridization patterns of marker genes from existing literature to estimate their spatial distribution (Siebert et al., 2019). However, due to the relatively small number of neurons sequenced (∼3,500) in that study, there was a possibility of missing some neuron populations. Additionally, we did not capture enough intermediate states of neuronal differentiation to fully resolve the transcriptional changes occurring as neurons develop from ISCs. Another study used single-cell SMART-seq2 to sequence 1,152 *H. vulgaris* interstitial cells, including 384 neurons (Klimovich et al., 2020). Although this method provided deeper sequencing, it did not substantially increase the number of neurons available for analysis.

We addressed previous limitations by sequencing 10-fold as many neurons, generating ∼35,000 single-cell transcriptomes of neurons and neural progenitors. This enhanced dataset allowed us to confirm the twelve neuron subtypes identified in our original study (Siebert et al., 2019) and to identify three previously unrecognized subtypes. Using fluorescent in situ hybridization (FISH), in situ hybridization chain reaction (HCR), and immunostaining, we refined the spatial map of the *H. vulgaris* nervous system, revealing how neuron subtypes are organized into established circuits characterized by shared neuropeptide expression. To further dissect the regulatory underpinnings of neural differentiation, we performed URD trajectory inference and ATAC-seq (Assay for Transposase-Accessible Chromatin using Sequencing) on sorted neurons to characterize the regulatory landscape during nervous system differentiation (Buenrostro et al., 2015; Farrell et al., 2018) and we demonstrate how these data can be used to formulate hypothetical gene regulatory networks (GRNs) for future testing. To support broad use and accelerate discovery in *H. vulgaris* neurobiology and regeneration, all data and tools are freely available through an interactive, user-friendly web portal.

## Results

### Expansion of the neuronal single cell transcriptome dataset for *Hydra vulgaris*

In our previous study, we used Drop-seq to create a single cell atlas of the adult *H. vulgaris* polyp, which included approximately 3,500 single-cell transcriptomes of differentiated neurons and cells undergoing neurogenesis (Macosko et al., 2015; Siebert et al., 2019). To build on this initial study, we aimed to increase the number of neural single-cell transcriptomes to uncover any additional molecular diversity in the *H. vulgaris* neuron repertoire that may have been missed due to the relatively small number of cells sequenced. To accomplish this, we performed scRNA-seq using the 10x Genomics platform to expand our dataset. To enrich for neurons and neural progenitors, we used Fluorescent Activated Cell Sorting (FACS) to collect cells from two different transgenic lines (**Fig. S1, Table S1**): (1) *Tg(actin1:GFP)*^rs3-in^, a previously published line in which GFP is expressed in all differentiated neurons, neural progenitors, and ISCs (Keramidioti et al., 2024), and (2) *Tg(tba1c:mNeonGreen)*^cj1-gt^, a line created for this study in which mNeonGreen is expressed under the control of the pan-neuronal gene *tba1c* and is therefore predicted to be expressed in all differentiated neurons (see Material and Methods for explanation of transgenic line naming scheme). However, we observed that certain neuron subtypes were underrepresented in the scRNA-seq data collected from this line, suggesting that our cloned promoter conferred some neuronal subtype specificity, possibly due to incomplete coverage of the *tba1c* regulatory region in our transgenic construct. (**Figs. S1, S2**).

We combined our new data with previously collected neuronal single-cell transcriptomes from our earlier study for downstream analysis (**Table S2**) (Siebert et al., 2019). Sequencing reads from all libraries were mapped to the *H. vulgaris* strain AEP gene models and processed following standard protocols (**Fig. S3**) (Cazet et al., 2023; Siebert et al., 2019). The expression of the pan-neuronal marker *elav2* was used to determine neuron identity and therefore remove non-neurons from the data set (Siebert et al., 2019). After filtering, we recovered 35,071 single-cell neural transcriptomes, expanding our dataset tenfold. We detected a median of 1371.5 genes and 2887.5 UMIs per cell (**Table S2**), a substantial improvement compared to the median of 563.5 genes and 1,082 UMIs per cell detected in the neuron-enriched Drop-seq libraries from our previous study (Siebert et al., 2019). To identify distinct neuron subtypes in our dataset, we used Seurat to perform Louvain clustering and visualized the results using Uniform Manifold Approximation and Projection (UMAP) (**Fig. 1B, Fig. S4, Table S3**) (Hao et al., 2021; McInnes et al., 2018; Satija et al., 2015; Stuart et al., 2019). Fifteen clusters of putative differentiated neurons were identified based on their high expression of *tba1c* and the absence of stem and progenitor gene expression (**Fig. S5**). As detailed below, all fifteen of these clusters were confirmed to consist of differentiated neurons. We performed non-negative matrix factorization (NMF) (Kotliar et al., 2019) to identify groups of co-expressed genes (“metagenes”), and we recovered metagenes for the fifteen neuron clusters, and stem and progenitor cells (**Fig. S6**).

### An integrated cell-type atlas of the *Hydra vulgaris* nervous system

This expanded scRNA-seq dataset provides a valuable resource for studying nervous system development, regeneration, and function in *H. vulgaris*. In this section, our goal is to present a comprehensive description of the *H. vulgaris* nervous system by integrating our new data with previously published work, including recent work by Keramidioti et al. (2024) using a novel pan-neural antibody (PNab) to provide a detailed anatomical description of the *H. vulgaris* nerve net. We begin by revisiting and expanding the neuron naming scheme originally proposed in Siebert et al., 2019, which defined eight neuron types further subdivided into subtypes. For each type/subtype, we describe roles in known neural circuits where applicable, determine neuropeptide expression patterns, and use FISH or HCR to characterize the spatial localization of subtypes, and in some cases identify novel subtypes (**Fig. 2**, **Fig. 3**). These integrated data provide a detailed classification of neuron subtypes within the *H. vulgaris* nervous system.

**Figure 2.**
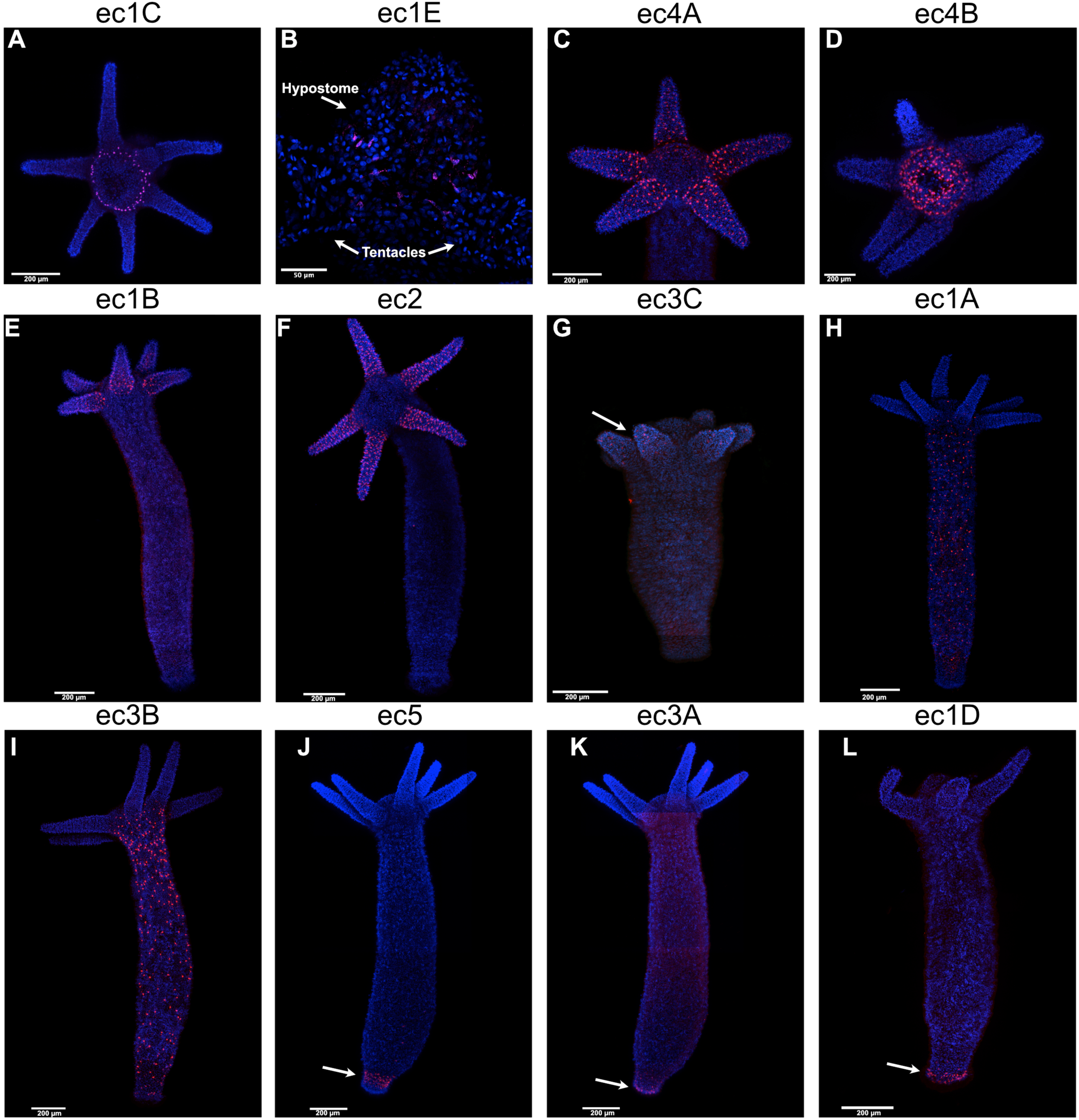
Spatial organization of ectodermal neuron subtypes in revealed by fluorescence in situ hybridization (FISH). Confocal images show FISH labeling of twelve distinct ectodermal neuron subtypes using subtype-specific molecular markers. **(A–G)** Oral neuron subtypes: ec1C (*G007791*), ec1E (*G003886*), ec4A (*RFamide C*, *G017226*), ec4B (*RFamide B*, *G017227*), ec1B (*G016169*), ec2 (*G010335*), and ec3C (*G026484*). Arrow in panel G indicates expression in the tentacles. **(H–I)** Body column neuron subtypes: ec1A (*G026993*) and ec3B (G004106). **(J–L)** Aboral neuron subtypes: ec5 (Hym176C, G016165), localized to the peduncle (arrow), and ec3A (G021930) and ec1D (G004200), localized to the basal disk (arrows). See Fig. S7 for whole animal images of ec4A, ec4B, and ec1C FISH experiments. Nuclei are counterstained with Hoechst (blue). All panels show standard FISH labeling except panel B, in which ec1E neurons required localization using hybridization chain reaction (HCR).

**Figure 3.**
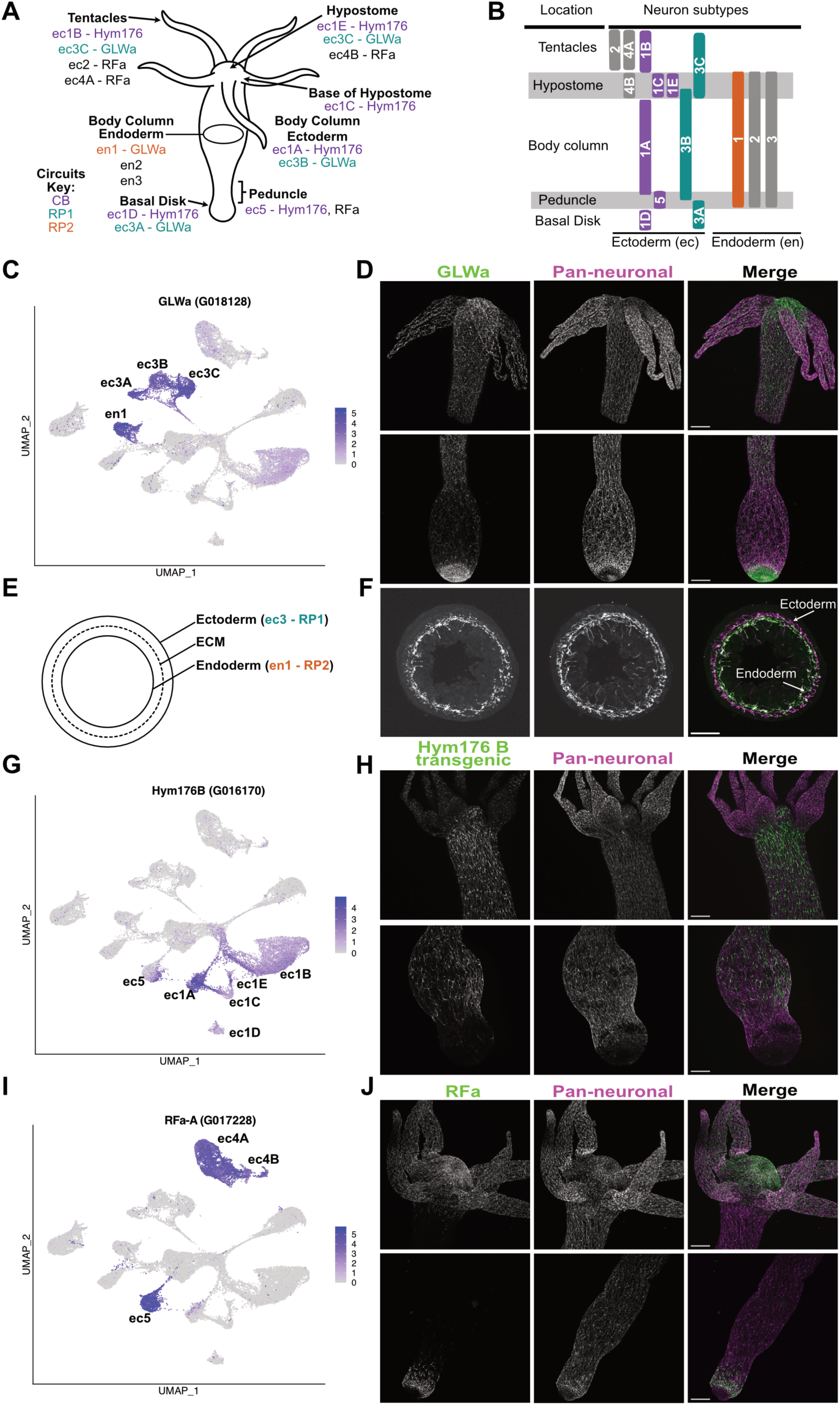
Neuropeptide expression delineates distinct neural circuits in *Hydra vulgaris*. **(A,B)** Schematics illustrate the spatial distribution of neuron subtypes along the *Hydra* body column. **(C)** The RP1 circuit comprises ec3A–C neurons, which express neuropeptides from the GLWa family. The RP2 circuit includes endodermal en1 neurons, which also express GLWa. **(D)** Immunostaining using an antibody against GLWa (green) reveals the RP1 circuit extending from the basal disk to the tentacles. **(E, F)** GLWa immunostaining in body column rings shows endodermal en1 neurons of the RP2 circuit. **(G)** The CB circuit includes ectodermal neuron subtypes ec1A–D and ec5, which express Hym176 neuropeptides. **(H)** GFP immunostaining (green) in the *Hym176B* reporter line marks CB circuit neurons along the body column. **(I)** The RFa-A neuropeptide is expressed in ec4 and ec5 neurons, the latter being part of the CB circuit. **(J)** Immunostaining with an antibody against RFa (green) reveals ec5 neurons in the peduncle and ec4 neurons near the oral end. In all immunostaining panels (D, F, H, J), co-labeling with PNab (magenta) highlights the full structure of the nervous system. Scale bars: 200 µm. The lower images in panel D and the upper images in panel H are reproduced from Keramidioti et al., 2024.

#### Classification scheme for H. vulgaris neurons

To systematically classify *H. vulgaris* neurons, we developed a hierarchical and expandable naming scheme that can accommodate future insights into neuronal function, morphology, and spatial organization. The *H. vulgaris* nervous system is comprised of two nerve nets, one embedded in the ectoderm and one in the endoderm (Keramidioti et al., 2024). The ectodermal nerve net extends from the basal disk to the tentacle tips, with higher neuron density at the oral (tentacle and hypostome) and aboral (peduncle and basal disk) regions as compared to the body column (Keramidioti et al., 2024). Along the length of the animal, ectodermal neurons project neurites that form bundles and communicate laterally through gap junctions (Keramidioti et al., 2024). Two major non-overlapping circuits, contraction burst (CB) and rhythmic potential 1 (RP1), run along the axis in the ectoderm and regulate behaviors such as longitudinal contractions and somersaulting (Dupre and Yuste, 2017; Yamamoto and Yuste, 2023). The endodermal nerve net is distributed along the body in the endodermal layer but is absent in the tentacles (Keramidioti et al., 2024). The rhythmic potential 2 (RP2) neural circuit, which is involved in feeding behavior, is part of the endodermal nerve net (Dupre and Yuste, 2017; Giez et al., 2023; Keramidioti et al., 2024; Siebert et al., 2019).

Building on our previous work, we first assigned neurons to either the ectodermal (“ec”) or endodermal (“en”) nerve nets, using expression of the endodermal marker gene *budhead* to guide this distinction (**Fig. 1D, Fig. S5**) (Siebert et al., 2019). Our prior classification grouped ectodermal neurons into five types (ec1-ec5); here, we confirm and extend this framework by showing that each type has a unique neuropeptide expression profile (**Fig. 1C, Table S4**). This neuropeptide-based strategy is consistent with approaches used to classify neuron types in *C. elegans*, mice, and humans (Das Gupta et al., 2021; Taylor et al., 2021; Zhong et al., 2022). As described in detail below, we further subdivided the five ectodermal types into twelve transcriptionally and spatially distinct subtypes (**Fig. 1C**, **Fig. 2**, **Fig. 3A,B**). In contrast, the endodermal nerve net is less complex, consisting of three transcriptionally distinct clusters that show moderate variation in neuropeptide expression (**Fig. 1C**). As described in detail below, these clusters correspond to three neuron types (en1-en3) (**Fig. 1C**). Overall, this classification extends the original ec1-ec5 and en1-en3 scheme described in Siebert et al. (2019) and provides a robust framework for future refinement as additional data emerge.

#### ec3 neurons

The RP1 circuit is composed of ec3 neurons, which express a suite of neuropeptides, most prominently *Hym-355* and products of the *GLWamide* gene (**Fig. 1C**) (Keramidioti et al., 2024; Leviev et al., 1997; Yamamoto and Yuste, 2023). To visualize the architecture of this circuit, we performed co-immunostaining using an antibody recognizing GLWamide peptides (Koizumi et al., 2015) and PNab (stains all neurons), revealing ec3 neurons extending throughout the length of the body column (**Fig. 3A-D**). ec3 neurons also uniquely express *innexin 6* and *innexin 14*, which likely contribute to gap junction formation and electrical coupling within the RP1 circuit (**Fig. S9B,C**) (Keramidioti et al., 2024). Consistent with our previously published data set, the ec3 neuron population forms a transcriptional continuum that can be divided into three clusters, each enriched for different marker genes **(Fig. 1B, Fig. S5**) (Siebert et al., 2019). We used fluorescent in situ hybridization (FISH) to map spatial distributions of these neuronal clusters and observed distinct localization along the oral-aboral axis, supporting classification as three ec3 subtypes. ec3A neurons are found in the basal disk, ec3B neurons are found in the body column and hypostome, and ec3C neurons are found in the tentacles (**Fig. 1B,D**, **Fig. 2G, I, K, Fig. 3A,B**).

#### ec1 and ec5 neurons

Together, ec1 and ec5 neurons comprise the CB circuit, which drives longitudinal contractions (Dupre and Yuste, 2017; Escobar et al., 2024; Noro et al., 2019; Noro et al., 2021). Both ec1 and ec5 neurons express *innexin-2*, a gap junction gene that facilitates communication between cells within the circuit (**Fig. S9A**) (Keramidioti et al., 2024; Siebert et al., 2019; Takaku et al., 2014). ec1 neurons express *FRamide* and members of the *Hym176* neuropeptide family, including *Hym176B* **(Fig. 1C**). We visualized ec1 neurons using a transgenic reporter line expressing GFP under the *Hym176B* promoter (Noro et al., 2019), combined with GFP and PNab co-immunostaining. This revealed the distribution of the ec1 neurons in the CB circuit, extending from just above the peduncle to the hypostome and tentacle base (**Fig. 3A,B,G,H)**. Co-immunostaining the *Hym176B* promoter line with PNab and GLWamide, a marker for ec3 neurons, confirmed that the body column ectodermal nerve net contains only ec1A and ec3B neurons (**Fig. 3A,B, Fig. S8**). ec5 neurons complete the CB circuit and share a similar neuropeptide expression profile with ec1, with additional expression of *Hym176C* and *RFamideA* (**Fig. 1C**). We confirmed their location in the peduncle using FISH for *Hym176C* (**Fig. 2J**) and RFamide immunostaining (**Fig. 3A,B,I,J**), noting that RFamide is also expressed in ec4 neurons (Koizumi et al., 2015).

In our previous study, we identified two distinct ec1 subtypes, ec1A and ec1B, based on distinct molecular markers (Siebert et al., 2019) (**Fig. S5**). Using FISH, we now confirm that ec1A neurons localize to the body column, whereas ec1B neurons are restricted to the tentacles, supporting their classification as separate subtypes (**Fig. 2E,H**). In our current dataset, we identified three additional clusters expressing *FRamide* and *Hym176* family neuropeptides, consistent with ec1 identity (**Fig. 1B, C**). FISH or HCR with cluster-specific marker genes confirmed three new neuron subtypes: ec1C, ec1D, and ec1E (**Fig. 2A,B,L, Fig. S5**). ec1C neurons form a ring around the mouth, potentially resembling the nerve ring previously described in *Hydra oligactis* (**Fig. 2A, Fig. S7C**) (Grimmelikhuijzen, 1985). ec1E neurons are located just above ec1C in the lower hypostome, while ec1D neurons localize to the basal disk (**Fig. 2B,L**).

Previous studies reported only a single neuronal population in the basal disk (Dupre and Yuste, 2017; Siebert et al., 2019). Therefore, to distinguish ec1D from ec3A neurons, we performed double FISH using subtype-specific markers, confirming their separation (**Fig. 4A-C**). Co-staining with GLWamide and PNab further supported the presence of a second, non-ec3A neuronal population in the basal disk, as we observed PNab-positive cells lacking GLWamide expression (**Fig. 4D-F**). In summary, the CB circuit is composed of six neuronal subtypes, ec5 and ec1A-E, which span the entire body axis from the basal disk (ec1D) to the tentacles (ec1B).

**Figure 4.**
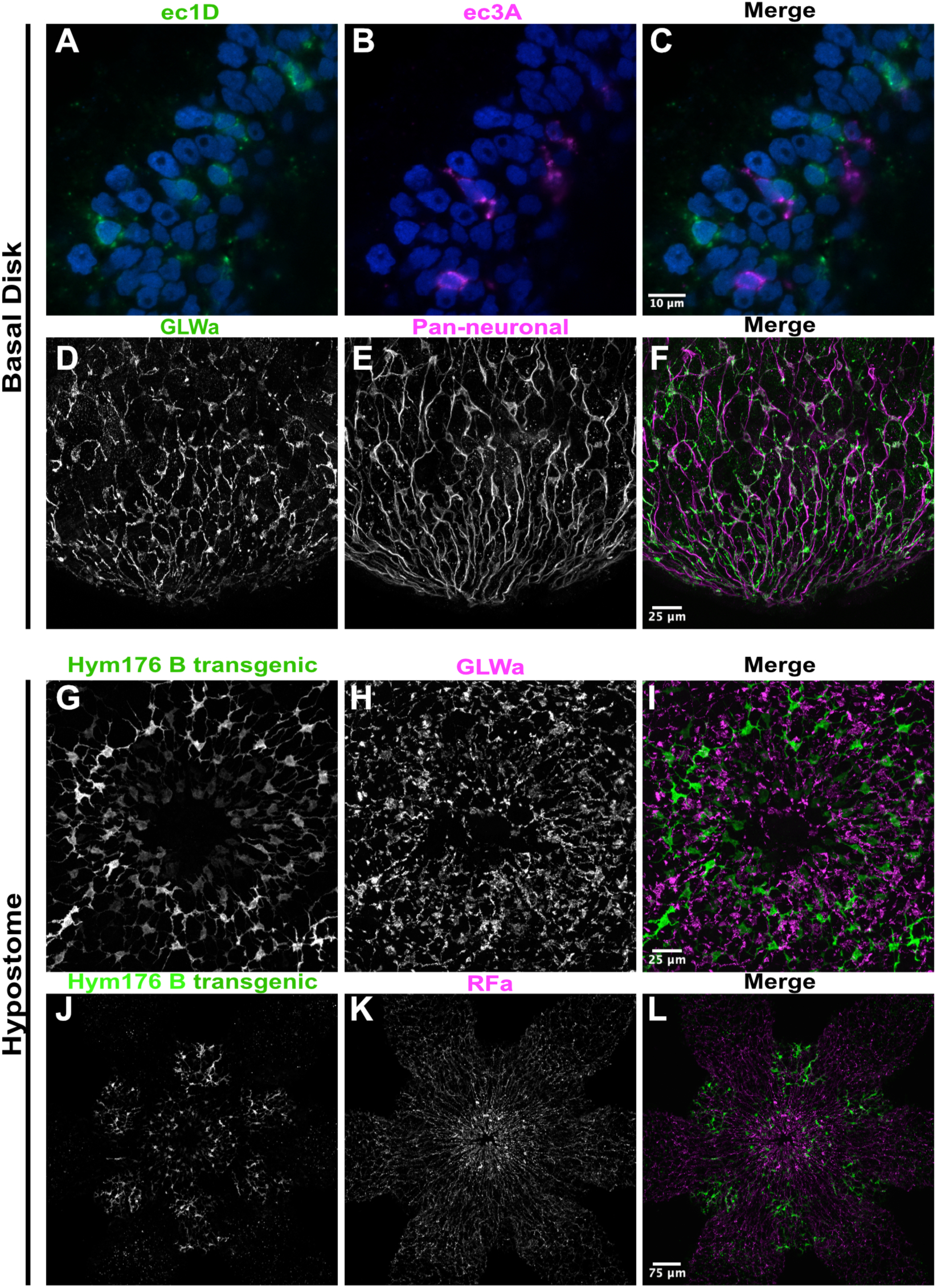
Localization of neuron subtypes in the basal disk and hypostome. **(A–C)** Double FISH using probes for *G021930* (magenta, ec3A) and *G004200* (green, ec1D) confirms that these neuron subtypes are distinct populations in the basal disk. Nuclei are counterstained with Hoechst (blue) **(D–F)** Immunofluorescence with antibodies against GLWa (green, ec3) and PNab (magenta, pan-neuronal) reveals GLWa-negative/PNab-positive neurons in the basal disk, consistent with ec1D identity. **(G–I)** Co-immunostaining in the *Hym17B-GFP* reporter line using anti-GFP (green) and anti-GLWa (magenta) shows the spatial relationship of ec1 and ec3 neurons in the hypostome. **(J–L)** Co-immunostaining in the Hym17B-GFP reporter line with anti-GFP (green) and anti-RFa (magenta) illustrates the positioning of ec1 and ec4B neurons in the hypostome.

For both the RP1 and CB circuits, the potential functional roles of the distinct neuronal subtypes remain unclear. However, differential gene expression within our dataset may provide insights into their specialization. Given the spatial restriction of these subtypes, observed gene expression differences may also reflect positional cues mediated by the Wnt signaling gradient originating from the oral end.

#### ec2 and ec4 neurons

In our previous work, we found that ec2 and ec4 neurons are localized to the oral end and likely do not participate in the RP1 or CB circuits (Keramidioti et al., 2024; Siebert et al., 2019). These neurons express distinct, non-overlapping sets of neuropeptide genes (**Fig. 1C)**. The ec4 cluster can be further subdivided into two transcriptionally distinct subtypes: ec4A and ec4B (**Fig. 1B, Fig. S5**). FISH analysis revealed that ec4A neurons are in the tentacles, while ec4B neurons are in the hypostome (**Fig. 2C,D, Fig. S8A,B**). Thus, four ectodermal neuron subtypes are in the hypostome (ec1C, ec1E, ec3B, and ec4B) and four are in the tentacles (ec1B, ec2, ec3C, and ec4A) (**Fig. 2**). The increased neuronal complexity at the oral end likely reflects the demands of coordinating tentacle and hypostome activity during behaviors such as feeding and locomotion.

We visualized the architecture of the hypostomal nerve net using immunostaining for RFamide (ec4), LWamide (ec3), and GFP in the *Hym176B* reporter line (ec1) (**Fig. 4G-L**). The tentacular nerve net remains less well resolved; however, ec2 neurons are likely sensory neurons embedded in the battery cell complex, an ectodermal epithelial cell that contains embedded nematocytes and the sensory neuron responsible for triggering nematocyte discharge (Hobmayer et al., 1990). In our whole-animal scRNA-seq dataset, ectodermal battery cells were sequenced along with their associated neurons, and we detected strong expression of the ec2 marker *RFamideC*, further supporting this interpretation (Siebert et al., 2019).

#### en1, en2, and en3 neurons

Compared to ectodermal neurons, endodermal neurons express fewer neuropeptide genes. en1 neurons express *GLWamide*, while en2 and en3 neurons have no strong neuropeptide expression (**Fig. 1C**). Immunostaining with the GLWamide antibody shows that en1 neurons form a ganglion-like network which constitutes the RP2 circuit (**Fig. 3A,E,F**). en1 neurons can communicate through gap junctions formed by *innexin 2* (**Fig. 9A**) (Dupre and Yuste, 2017; Keramidioti et al., 2024; Siebert et al., 2019). In our previous study, we identified en2 neurons as sensory cells using a transgenic reporter line; these neurons are characterized by a short apical sensory cilium projecting into the gastric cavity and a longer basal neurite that connects with the en1 neuron network (Keramidioti et al., 2024; Siebert et al., 2019). en2 cells express *innexin 15* (**Fig. S9 D**) and could form gap junctions but do not appear to form a neural circuit. To investigate the identity of en3 neurons, we generated a transgenic line expressing mNeonGreen under the control of the en3-specific gene *G022927* regulatory region. Confocal imaging of this reporter line revealed that en3 neurons are an additional type of sensory neuron (**Fig. S10**). Notably, a scanning electron microscopy study from four decades ago described two types of endodermal sensory neurons with unique morphologies, likely corresponding to the en2 and en3 neuron types in our data set (Westfall and Epp, 1985). Taken together with their different transcriptional profiles, these morphological differences support the classification of en2 and en3 as separate neuron types.

### Developmental trajectory inference of the homeostatic *H. vulgaris* nervous system

The *H. vulgaris* nervous system undergoes continuous renewal, with complete neuronal turnover approximately every three weeks (Bode et al., 1988). As a result, scRNA-seq of homeostatic adult *H. vulgaris* captures a dynamic mixture of stem cells, differentiated neurons, and cells in various stages of differentiation (Siebert et al., 2019). Given the increased number of neuronal progenitors captured in our new data set which provide information about the developmental relationships between neuronal cell types, we reconstructed developmental trajectories under steady-state conditions to describe the molecular events as these diverse neuron types are specified and differentiate. To achieve this, we applied URD, a diffusion-based algorithm that infers developmental trajectories as branching trees (**Fig. 5A, Fig. S11**) (Farrell et al., 2018), using ISCs as the root, defined by the marker *Hy-icell1* (G002332) (Siebert et al., 2019), and terminal tips defined by neuron subtype-specific markers and absence of progenitor markers (**Fig. S5**). Our analysis predicted the relationship between ISCs and eleven terminal branches of neurons. Limitations of the analysis are that ec4A and ec4B were collapsed into a single group, as trajectory connections to ISCs could not be resolved separately, and the newly described ec1C-E subtypes were excluded because progenitors that link them to ISCs were seemingly not captured.

**Figure 5.**
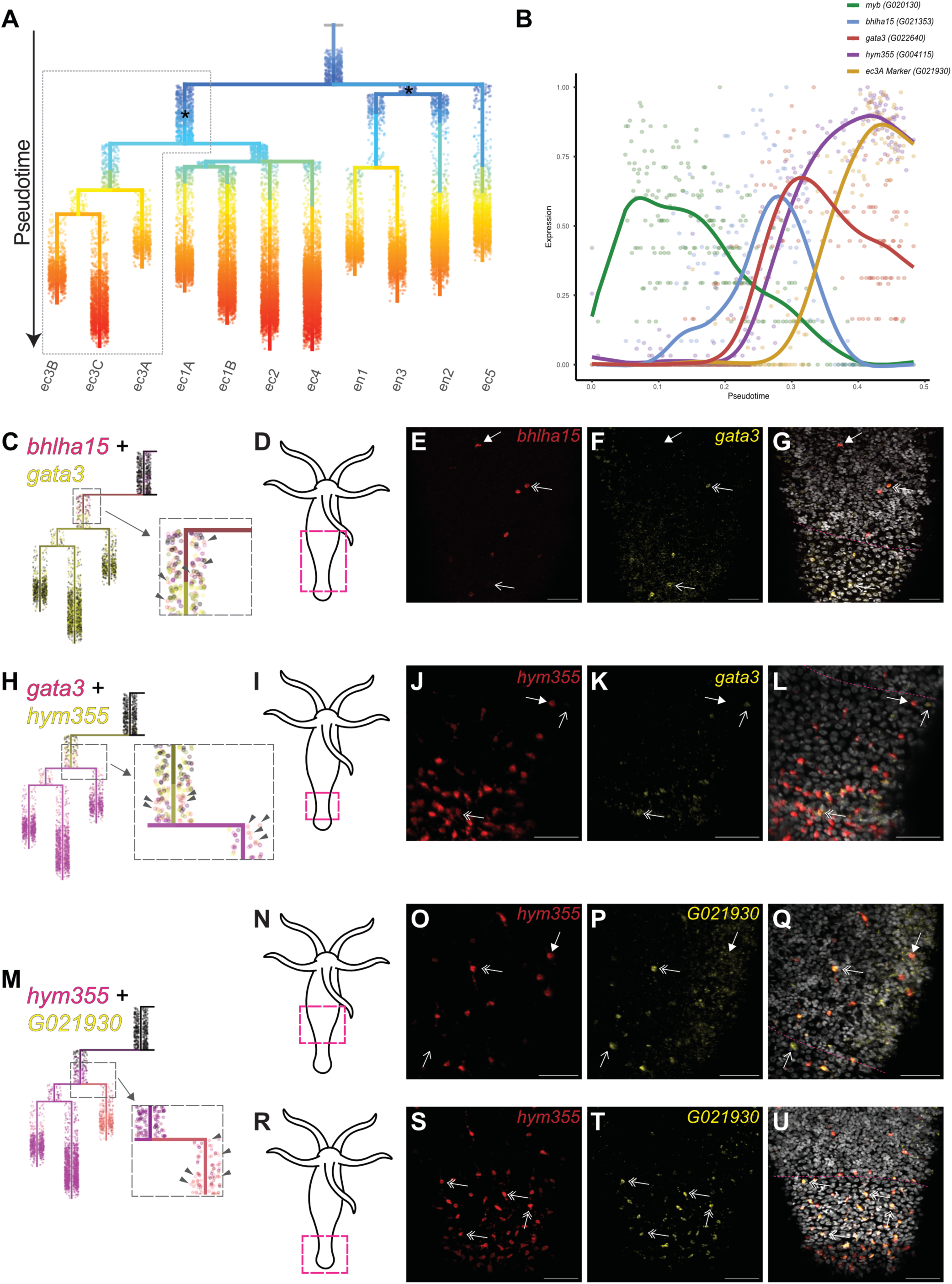
Trajectory reconstruction of neurogenesis reveals distinct developmental pathways for ectodermal and endodermal neurons. **(A)** Differentiation trajectories for eleven *Hydra* neuron subtypes were reconstructed using URD from single-cell transcriptomics data (Farrell et al., 2018). Interstitial stem cells (ISCs) were set as the “root” or starting point, and each neuron subtype served as a “tip” or endpoint. The tree is colored by pseudotime (developmental progression), with earlier pseudotime at the top (blue) and later pseudotime at the bottom (red). The boxed region highlights areas detailed in panels C, H, and M. Asterisks indicate segments corresponding to early differentiation events from ISCs into ectodermal or endodermal neuronal progenitors. **(B)** Spline plot showing expression trends for five genes during ec3A differentiation. The x-axis represents pseudotime (earlier on the left, later on the right), and the y-axis represents scaled gene expression levels. Each point is the average expression for five cells. **(C-U)** Double FISH validates predicted transition states during ec3A differentiation. Expression states in FISH images are indicated by arrow types: closed arrows for cells expressing only the first gene, open arrows for cells expressing only the second, and double arrows for cells co-expressing both genes. **(C-G)** Validation of early ec3A transition states co-expressing *bhlha15* and *gata3*. **(C)** Visualization of *bhlha15* and *gata3* expression on the URD trajectory. Co-expressing cells (orange) are indicated with arrowheads; black represents cells expressing neither gene. **(D-G)** Confocal microscopy of double FISH for early transitions: **(D)** Body region imaged, (E) *bhlha15* expression (red), (F) *gata3* expression (yellow), and **(G)** Overlay with Hoechst-stained nuclei (gray). **(H-L)** Validation of mid ec3A transition states co-expressing *hym355* (magenta) and *gata3* (yellow). **(H)** Visualization of *hym355* and *gata3* expression on the URD trajectory. Co-expressing cells (orange) are highlighted with arrowheads. **(I-L)** Confocal microscopy of double FISH for mid transitions: **(I)** Body region imaged, (J) *hym355* expression (red), (K) *gata3* expression (yellow), and **(L)** Overlay with Hoechst-stained nuclei (gray). **(M-U)** Validation of late ec3A transition states co-expressing *hym355* (magenta) and ec3A marker *G021930* (yellow). **(M)** Visualization of *hym355* and *G021930* expression on the URD trajectory. Co-expressing cells (orange) are marked with arrowheads. **(N-U)** Confocal microscopy of double FISH for late transitions: **(N, R)** Body region imaged, (O, S) *hym355* expression (red), (P, T) *G021930* expression (yellow), and **(Q, U)** Overlay with Hoechst-stained nuclei (gray). Scale bars: 50 µm. Pink dotted lines in microscopy images (G, L, Q, U) indicate the boundary between the body column and peduncle, determined by nuclei morphology.

We previously showed that ISCs transitioning to neuronal fates express the transcription factors *myc3* and *myb*, which are downregulated upon terminal differentiation (Siebert et al., 2019). Here, our trajectory analysis identified three early-branching populations expressing both *myc3* and *myb* (**Fig. S12**). Two populations are multi-potent progenitors, one of which gives rise to endodermal neurons (en1, en2, en3), and the other of which gives rise to most ectodermal neuron types (ec1–ec4) (**Fig. 5A**). Notably, ec5 neurons appear to differentiate directly from ISCs, though this may reflect incomplete capture of intermediate states (**Fig. 5A**). Together, these findings suggest that endodermal and ectodermal neurons each arise from distinct progenitor pools that become progressively fate-restricted during differentiation, ultimately generating the full diversity of *H. vulgaris* neuron subtypes.

The progressive specification and differentiation of neuronal subtypes predicted here will be accompanied by a cascade of transcriptional events that regulate each step. As proof of principle, we used our trajectory analysis to visualize the expression dynamics of transcription factors and terminal marker genes that belong to one such cascade during ec3A differentiation. Plotting the expression of *myb*, *bhlha15*, *gata3, hym355*, and ec3A marker *G021930* shows their sequential expression accompanied by transition states with temporary co-expression (**Fig. 5B**). To validate these predicted transition states, we performed double fluorescent *in situ* hybridization (FISH) using pairs of genes with overlapping expression domains across three predicted transition states (**Fig. 5C-U**). For gene pair, *bhlha15* and *gata3* (**Fig. 5C-G**), *gata3* and *hym355* (**Fig. 5H-L**), and *hym355* and *G021930* **(Fig. 5M-Q**), we identified cells expressing each gene individually as well as cells co-expressing both genes, consistent with our predicted differentiation trajectory.

Notably, most neurons co-express the terminal marker genes *hym355* and *G021930* in the basal disk (**Fig. 5M, R-U**), which aligns with the expected localization of terminally differentiated ec3A neurons (**Fig. 2K**). These findings validate our trajectory analysis and demonstrate its ability to accurately identify gene expression dynamics during the differentiation of neuronal subtypes in *H*. *vulgaris*. This trajectory analysis is available as part of our online portal to enable visualization of similar gene expression cascades during the specification of each *H*. *vulgaris* neuronal subtype.

### Characterizing the chromatin landscape in the *H. vulgaris* nervous system

Our work establishes *H. vulgaris* as one of the few adult organisms in which the nervous system has been transcriptionally defined, including a trajectory analysis describing the molecular changes that occur during neuronal differentiation. This provides a unique opportunity to identify the GRNs that describe the differentiation of all neurons in an adult nervous system. To advance this goal, we analyzed the expression of transcription factors (TFs) in our data set. We previously used phylogenetic analysis to identify 811 putative TFs, defined as genes with predicted DNA binding domains, with detectable expression in *H. vulgaris* (Cazet et al., 2023). We identified the expression patterns of these TFs across our neuronal cell types, finding that 665 of those 811 TFs are expressed within the nervous system, and we used weighted gene co-expression network analysis (WGCNA) to cluster them based on average expression in each cell type (**Fig. S13**) (Langfelder and Horvath, 2008). Most neuronal subtypes are characterized by the unique expression of at least a dozen TFs, and most broader classes of neurons (e.g. ec1A/B/C/D/E) are also characterized by a few TFs with shared expression across the class.

To build GRNs underlying nervous system development in *H. vulgaris*, it is essential to identify the cis-regulatory elements (CREs) bound by neuronal transcription factors. As a foundational step towards this goal, we performed ATAC-seq on FACS-isolated neurons from the *Tg(actin1:GFP)*^rs3-in^ and *Tg(tba1c:mNeonGreen)*^cj1-gt^ transgenic lines (**Table S5**). These neuron-specific datasets were of high quality and showed strong replicate concordance, meeting ENCODE standards (**Fig. S14, Table S6**) (Landt et al., 2012). Importantly, the data quality matched that of our previously published whole-animal ATAC-seq from *H. vulgaris*, indicating that FACS isolation did not compromise chromatin integrity (Cazet et al., 2023). We observed significantly increased accessibility near neuronal genes in the neuron-specific libraries compared to whole-animal data (p < 0.001 for each line; **Fig. 6A, B**). Moreover, this approach uncovered accessible regions at neuronal loci that were not detected in whole-animal, while confirming a lack of accessibility at non-neuronal genes such as *wnt3* (**Fig. 6C-H**) (Cazet et al., 2023). These findings provide an essential resource for defining the regulatory architecture of *H. vulgaris* neurogenesis.

**Figure 6.**
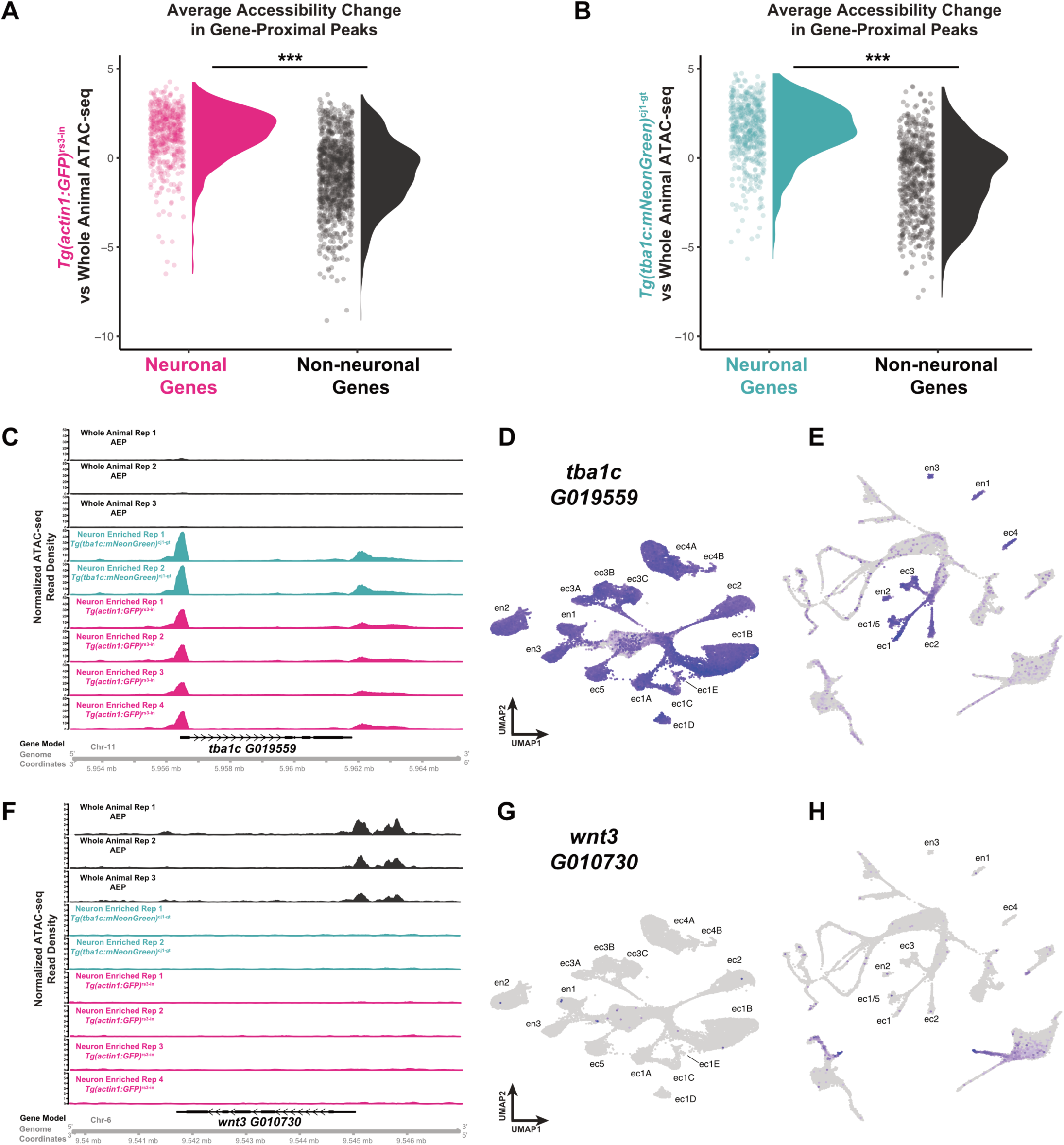
Characterization of the neuronal chromatin landscape using ATAC-seq. (**A-B**) Neuron-enriched ATAC-seq libraries were generated from six samples: two from the *Tg(tba1c:mNeonGreen)^cj1-gt^* transgenic line (green) and four from the *Tg(actin1:GFP)^rs3-in^* transgenic line (pink) (Keramidioti et al., 2024). To assess neuronal peak enrichment, three previously published whole-animal ATAC-seq libraries (AEP1-3; Cazet et al., 2023) were used as a reference. Panels (**A**) and (**B**) show the average accessibility changes in gene-proximal peaks for *Tg(actin1:GFP)^rs3-in^*and *Tg(tba1c:mNeonGreen)^cj1-gt^*, respectively, compared to whole-animal data sets. Neuronal genes exhibit significant peak enrichment within 10,000 base pairs of the transcription start site relative to non-neuronal genes (p < 0.001, t-test). (**C**) ATAC-seq tracks for the *alpha tubulin* locus (*tba1c*, *G019559*) show peaks in the neuron-enriched libraries that are absent in the whole animal libraries. (**D**) UMAP visualization of the neuron-enriched scRNA-seq dataset shows that *tba1c* is ubiquitously expressed in neurons. (**E**) UMAP visualization of the whole-animal scRNA-seq dataset shows that *tba1c* expression is restricted to neurons (Cazet et al., 2023; Siebert et al., 2019). (**F**) ATAC-seq tracks for the *wnt3* locus (*G010730*) show peaks in the whole-animal libraries that are absent in the neuron-enriched libraries. (**G**) UMAP visualization of the neuron-enriched scRNA-seq dataset shows no expression of *wnt3* in neurons. (**H**) UMAP visualization of the whole animal scRNA-seq dataset shows that *wnt3* is expressed in the oral ectodermal and endodermal epithelium (Cazet et al., 2023; Siebert et al., 2019).

### Gene regulatory network inference demonstrated with ec4 neuron specification

To illustrate the utility of the data presented in this study for hypothesis generation, we constructed a putative GRN describing the differentiation of the oral neuron type ec4 (**Fig. 7**). Reasoning that oral neurons might share core regulatory features, we first identified transcription factors expressed during the differentiation of ec1B, ec2, ec3C, and ec4 neurons, and visualized their expression dynamics along pseudotime using spline plots (**Fig. 7A, Fig. S15A**). As observed previously, *Myb* and *Myc3* are activated early during neurogenesis across all neuronal subtypes, marking the onset of neuronal specification (Siebert et al., 2019). Here we show that *Sox2* is expressed broadly across oral neuron subtypes (**Fig. 7B, Fig. S15B**) and that open chromatin near the *Sox2* locus has both *Myb* and *Tcf* binding sites, consistent with the possibility that Sox2 expression within oral neurons is directly regulated by *Myb* and receives inputs from the Wnt signals emanating from the oral organizer (**Fig. 7H, Fig. S15C**).

**Figure 7.**
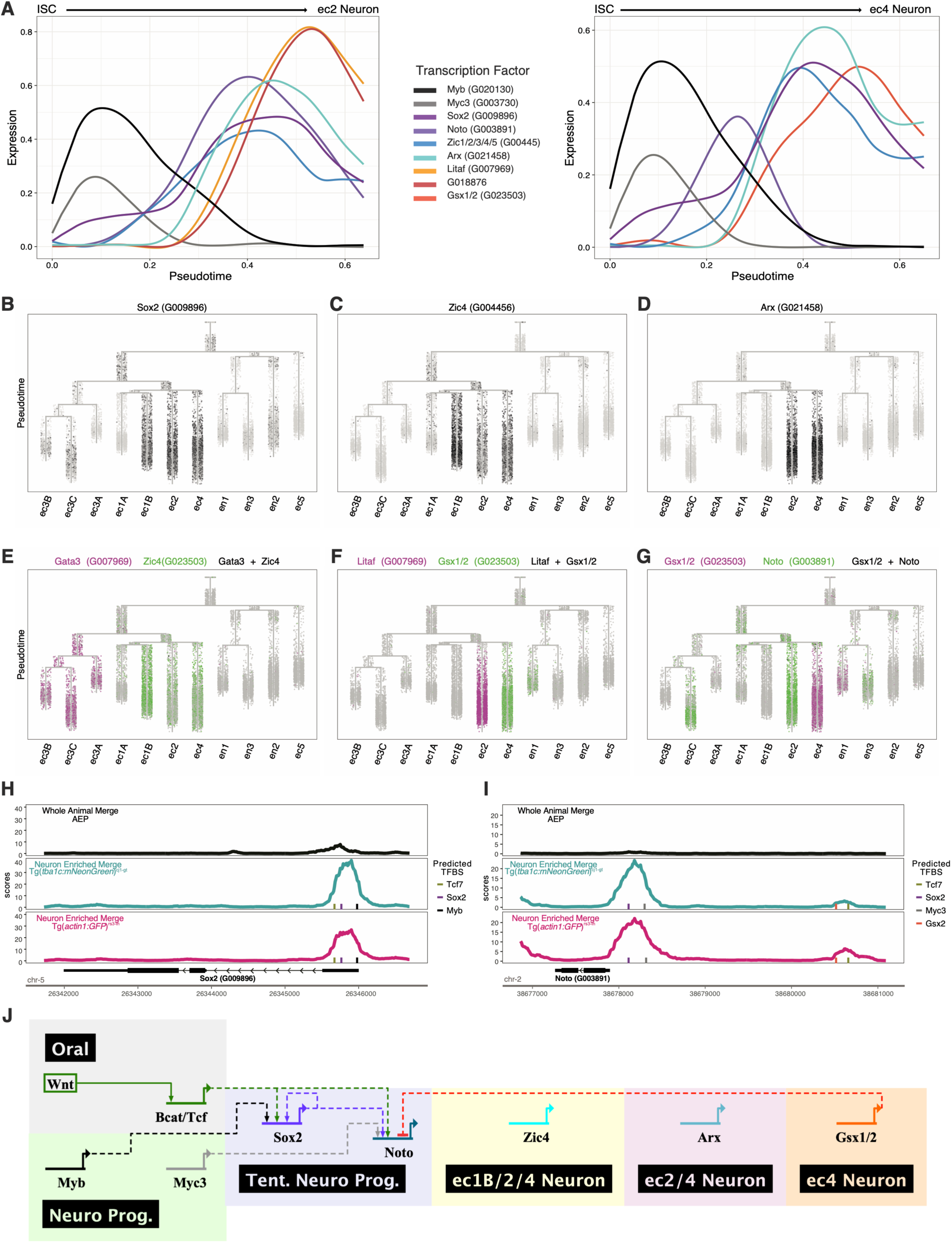
A putative gene regulatory network (GRN) underlying ec4 neuron specification. **(A)** Spline plots showing transcription factor expression dynamics during ec2 and ec4 differentiation. **(B)** *Sox2* (*G009896*) is broadly expressed in oral neuron subtypes ec3C, ec1B, ec2, and ec4. **(C)** *Zic4* (*G004456*) is expressed in ec1B, ec2, and ec4. **(D)** *Arx* (*G021458*) is restricted to ec2 and ec4. **(E)** Co-expression trajectory tree of *Gata3* (*G022640*, magenta) and *Zic4* (*G004456*, green) show mutually exclusive expression. **(F)** Co-expression trajectory tree of *Litaf* (*G007969*, magenta) and *Gsx1/2* (*G023503*, green) reveals subtype-specific expression in ec2 and ec4 neurons. **(G)** Co-expression trajectory tree of *Gsx1/2* (magenta) and *Noto* (G003891, green) shows mutually exclusive expression in ec4. **(H)** ATAC-seq tracks at the *Sox2* locus reveal open chromatin regions containing *Tcf*, *Sox2*, and *Myb* binding motifs. (I) ATAC-seq tracks at the Noto locus highlight peaks containing *Tcf*, *Sox2*, *Myc*, and *Gsx2* binding motifs. (J) A proposed GRN model summarizing transcriptional regulators and regulatory relationships involved in ec4 neuron specification in *Hydra vulgaris*. Plots in this figure are made using the scRNA-seq portal (https://research.nhgri.nih.gov/HydraAEP/SingleCellBrowser/).

As differentiation progresses, the oral neurons separate into two groups: ec1B/ec2/ec4, which express *Zic4*, and ec3C, which express *Gata3* (**Fig. 7C,E, Fig. S15D,F**). Previous studies have described antagonism between *Zic4* and *Gata3* in the ectodermal epithelium, a similar mutual antagonism may regulate this cell fate decision during neurogenesis (Ferenc et al., 2024; Vogg et al., 2022). Following *Zic4* expression, the ec4 neuronal identity becomes increasingly refined, first when *Arx* is jointly expressed in both ec2 and ec4, which is subsequently followed by the differential expression of *Litaf* in ec2 and *Gsx1/2* in ec4 (**Fig. 7D,F, Fig. S15E,G**).

Although the mechanism driving *Gsx1/2* expression specifically in ec4 neurons is not yet clear, our data suggest a regulatory interaction between *Gsx1/2* and the transcription factor *Noto* that may be key to stabilizing ec4 fate. *Noto* is expressed relatively early in the differentiation of oral neurons, and accessible binding sites for both *Myc3* and *Tcf* suggest possible upstream inputs (**Fig. 7A,G,I**). Interestingly, *Noto* expression declines in ec4 neurons as *Gsx1/2* expression increases (**Fig. 7A,G, Fig. S15H**). Given that the *Noto* locus contains an accessible *Gsx2*-binding site (**Fig. 7I**), and that *Gsx1/2* TFs can function as repressors when acting as monomers (Salomone et al., 2021), we propose that *Gsx1/2* represses *Noto* in ec4 neurons to reinforce ec4 identity. In contrast, *Noto* remains expressed in differentiated ec2 and ec3C neurons (**Fig. 7F,G, Fig. S15I**). Based on the observations described in this section, we constructed a testable GRN model for ec4 neuron specification (**Fig. 7J**).

### A broad community resource

Our results for ec4 neurons support the value of the data presented here for generating testable hypotheses to understand neuronal development in *Hydra vulgaris*. To support on-going and future research in *Hydra* neuroscience, we have made our datasets accessible through interactive web portals. Our scRNA-seq data can be explored via a user-friendly browser (https://research.nhgri.nih.gov/HydraAEP/SingleCellBrowser/), which allows researchers to visualize gene expression across neuron types using UMAP projections, dot plots, and trajectory branching trees and spine plots; examples of plots derived from our portal are shown in figures 7 and S15. The portal also includes functional annotations and conversion tools for matching gene IDs across *H. vulgaris* genomic and transcriptomic references. In parallel, we have integrated our neuron-specific ATAC-seq data into our existing *H. vulgaris* AEP genome browser (https://research.nhgri.nih.gov/HydraAEP/), enabling visualization of chromatin accessibility near neural genes. Easy access to our portals is provided through the community Resource Hub (openhydra.org), which offers centralized links to these high-resolution datasets. These tools are designed to facilitate broad data accessibility and accelerate discovery in *Hydra* neuroscience.

## Discussion

The number of research organisms used in neuroscience has grown significantly in recent years, driven by advances in single cell transcriptomics that enable the exploration of neuronal diversity across a broad range of species, including less conventional research organisms (Chari et al., 2021; Cole et al., 2024; Fincher et al., 2018; Hulett et al., 2024; Sebé-Pedrós et al., 2018a; Sebé-Pedrós et al., 2018b). Despite this progress, comprehensive transcriptional characterizations of adult nervous systems remain limited. Here, we present a detailed molecular map of the adult *H. vulgaris* nervous system based on ∼35,000 single-cell transcriptomes, including differentiated neurons and cells undergoing neurogenesis. In addition to confirming the twelve neuron subtypes described previously (Siebert et al., 2019), we identified three novel subtypes in the ectodermal nerve net. For each ectodermal neuron subtype, we defined unique cell markers and determined their spatial distribution, providing the first high-resolution map of the *H. vulgaris* ectodermal nerve net. Our study also confirmed the presence of the three previously described endodermal neuron subtypes (Siebert et al., 2019) and we identify the en3 subtype as having sensory neuron morphology. We propose a nomenclature scheme for *H. vulgaris* neuronal subtypes, designed to evolve with future discoveries.

*H. vulgaris* offers numerous experimental advantages, including small size, optical transparency, and accessible transgenic tools, which have accelerated efforts to link neural circuits to behavior. In a landmark study, the first use of GCaMP in *H. vulgaris* revealed three distinct longitudinal circuits (CB, RP1, RP2) that coordinate basic behaviors (Dupre and Yuste, 2017). Building on this work, others have linked specific neuron types and circuits to behaviors such as somersaulting and feeding (Giez et al., 2023; Yamamoto and Yuste, 2023), while innovative tools now allow precise sensory stimulation and automated behavior tracking (Badhiwala et al., 2018; Badhiwala et al., 2021; Escobar et al., 2024; Gonzales et al., 2017; Kim et al., 2024; Tzouanas et al., 2021). The transcriptional and spatial map presented here complements and expands this functional toolkit. By identifying subtype-specific markers and chromatin accessibility patterns, our dataset will facilitate targeted genetic manipulation and help to uncover the molecular mechanisms underlying neuronal function.

*H. vulgaris* is also a unique model for nervous system development and regeneration. In uninjured adults, ISCs continuously replace neurons, supporting a dynamic state of homeostasis. ISCs also enable complete nervous system regeneration following catastrophic injury, with a single ISC capable of regenerating the entire nervous system (David and Murphy, 1977). These features provide an opportunity to compare homeostatic and regenerative neurogenesis within a single organism and uncover regulatory mechanisms specific to either process. Using trajectory inference, we mapped the gene expression programs driving ISC differentiation into neuron subtypes under steady-state conditions, establishing a foundation for future comparisons with regeneration. Future research should aim to connect the molecular events driving neuronal differentiation with the acquisition of neural circuit synchronization and the recovery of behaviors during regeneration. In recent work, we correlated the timing of ec5 neuron differentiation with re-synchronization of the CB circuit during behavioral recovery, highlighting the value of integrating molecular and functional readouts (Escobar et al., 2024). This approach will be essential for understanding how circuits are reassembled and behaviors restored after injury.

In summary, we provide an unparalleled molecular characterization of the *H. vulgaris* nervous system, including newly identified neuron subtypes, a spatial map of ectodermal neurons, and a framework for building GRNs to study neuronal differentiation. These resources advance *H. vulgaris* as a powerful model for nervous system biology and are publicly available through user-friendly web portals to support broad use by the research community and inspire new investigators to explore this regenerative system.

## Materials and Methods

### *Hydra vulgaris* Strains and Culturing Conditions

All *Hydra vulgaris* strains used in this study were cultured in *Hydra* medium (0.38mM CaCl_2_, 0.32 mM MgSO_4_ X 7H2O, 0.5 mM NaHCO_3_, 0.08 mM K_2_CO_3_) at 18°C with a 12 hr light-dark cycle. *Hydra* were fed freshly hatched, premium grade *Artemia nauplii* (https://www.brineshrimpdirect.com) 2-4 times a week and were transferred into fresh *Hydra* medium 4-8 hours after feeding. Animals were starved for at least 48 hours prior to experimentation.

### Generation of transgenic strains

Transgenic *Hydra* lines *Tg(tba1c:mNeonGreen)^cj1-gt^*and *Tg(G022927:mNeonGreen)^cj1-i^* were generated using established protocols (Juliano et al., 2014b; Klimovich et al., 2019; Wittlieb et al., 2006). Promoter regions of 1901 bp (*tba1c*) and 1576 bp (*G022927*) upstream of their respective transcription start sites were PCR-amplified from *Hydra vulgaris* (strain 105) genomic DNA using Phusion™ High-Fidelity DNA Polymerase (ThermoFisher Scientific, #F530S), with a 55°C annealing and 60°C extension temperature. BamH1 and Xba1 restriction sites were added via the forward and reverse primers for cloning into the reporter plasmid upstream of mNeonGreen (**Table S7**).

PCR products were gel-purified, digested at 37°C for 10 minutes using BamH1 and Xba1 FastDigest enzymes (ThermoFisher Scientific, #FD0054 and #FD0684), and purified with the Zymo DNA Clean & Concentrator Kit (#D4003). Digested fragments were ligated into the BsAmp_mNG_tdT_backbone plasmid using Promega 10X Fast Ligase buffer (#M1801) for 4 hours at room temperature. Ligation products were transformed into DH5α-competent E. coli, followed by plasmid miniprep (Qiagen, #27106) and Sanger sequencing for insert verification. Final plasmids were amplified via maxiprep (Qiagen, #12162) and eluted in autoclaved Milli-Q water for microinjection.

Microinjections were performed into 1-cell stage *Hydra vulgaris* AEP embryos as previously described (Juliano et al., 2014b), with the following modifications: (1) the injection solution consisted of 1 µL of 0.5% phenol red (Sigma #P0290-100ML) mixed with 6 µL of plasmid DNA, followed by centrifugation; (2) embryos were fertilized for 1–2 hours before injection. Embryos were injected using an Eppendorf FemtoJet 4x and InjectMan NI 2 microinjector under a Leica M165 C stereo microscope. Transgenic animals were propagated asexually. Germline transmission was confirmed by crossing male and female Tg(tba1c:mNeonGreen)cj1-in animals to produce an F1 population.

### Naming scheme for transgenic lines

We have established a standardized naming convention modeled after the zebrafish community guidelines to ensure that each transgenic Hydra line has a unique and informative identifier. The format is: **Tg(promoter:gene_or_reporter)^lab#-lineage**, where **Tg** designates the line as transgenic. The **promoter** indicates the regulatory element driving expression, and **gene_or_reporter** specifies the transgene product (e.g., GFP or a specific gene). The **lab** field uses the initials of the principal investigator to denote the lab of origin (e.g., “cj” for the Juliano lab, “rs” for the Steele lab), followed by a unique allele number to distinguish independent integration events. The **lineage** suffix indicates the cell lineage in which the transgene is integrated—**ec** for ectoderm, **en** for endoderm, **in** for interstitial—or **gt** if the transgene has been germline transmitted. Using this naming scheme, we have renamed the transgenic line previously described in both Siebert et al., 2019 and Keramidioti et al., 2024 as *Tg(actin1:GFP)^rs3-in^*. In this line, GFP is driven by the actin1 promoter and is integrated into the interstitial lineage. However, likely due to positional effects of the integration site, GFP expression is largely restricted to neurons and neuronal progenitors.

### Dissociation of *Hydra* into Single Cells for Fluorescence-Activated Cell Sorting (FACS)

Forty to forty-five asexual, starved *Hydra* polyps were washed three times in sterile *Hydra* medium and transferred to 1.5 mL Eppendorf tubes. Medium was replaced with ∼75 U/mL Pronase E (VWR, E629-1G) in 1 mL of room temperature Dissociation Medium (DM; 5 mM CaCl₂·2H₂O, 1 mM MgSO₄·7H₂O, 2.8 mM KCl, 11 mM HEPES, 0.67 mM Na₂HPO₄, 0.44 mM KH₂PO₄, 5 mM sodium pyruvate, and 5 mM sodium citrate·2H₂O; pH 6.9–7.0). Polyps were dissociated for 90 minutes at 22–24°C with gentle agitation. Dissociated cells were transferred to a small Petri dish in 2 mL DM, gently pipetted up and down 5–10 times using a 1000 µL pipette, and filtered through a pre-soaked 70 µm Corning® strainer (Sigma #CLS431751) into a 50 mL conical tube containing 1 mL DM. The Petri dish and strainer were rinsed with an additional 1 mL DM. Cells were centrifuged at 300 g for 5 minutes (brake off), resuspended in 2 mL DM, and re-spun under the same conditions. The pellet was resuspended in 1 mL DM and filtered through a pre-soaked 40 µm Corning® strainer (Sigma #CLS431750) into a fresh conical tube.

### Fluorescence-Activated Cell Sorting (FACS)

FACS was used to collect fluorescent cells from transgenic *Hydra* lines *Tg*(*tba1c:mNeonGreen*)⁽ᶜʲ¹⁻ᶠ¹⁾ and Tg(*actin1:GFP*)⁽ʳˢ³⁻ⁱⁿ⁾ for both scRNA-seq and ATAC-seq. For scRNA-seq, one library was prepared from bud-free *Tg*(*tba1c:mNeonGreen*) ⁽ᶜʲ¹⁻ᶠ¹⁾ animals, and three libraries from *Tg*(*actin1:GFP*) animals, one from bud-free and two from budding animals. For ATAC-seq, two libraries were generated from bud-free *Tg*(*tba1c:mNeonGreen*) ⁽ᶜʲ¹⁻ᶠ¹⁾ animals and four from non-budding *Tg*(*actin1:GFP*)⁽ʳˢ³⁻ⁱⁿ⁾ animals. In all cases, fluorescent cells were sorted using a 100 µm nozzle on a MoFlo Astrios EQ Cell Sorter (Beckman Coulter) into 400 µL dissociation medium (DM) supplemented with 0.01% BSA. For ATAC-seq, sorting was performed into DNA LoBind® tubes (Eppendorf #0030122348), and 1 mL of additional DM was added prior to sorting to bring the final volume to 2 mL. Cells were pelleted at 300 g for 5 minutes and resuspended in 50 µL DM + 0.01% BSA for scRNA-seq, or counted using Hoechst staining and a Fuchs-Rosenthal hemocytometer (InCyto DHC-F01-5) for ATAC-seq. FACS gating used *Tg*(*actin1:GFP*)⁽ʳˢ³⁻ⁱⁿ⁾ and *Hydra vulgaris* AEP as positive and negative controls, respectively, and followed Siebert et al. (2019).

### 10x Genomics Single Cell RNA Library Preparation, Sequencing, and Analysis

A step-by-step description of the scRNA-seq cluster analysis, including all relevant code, is provided in the markdown document *01_hydraNeuronAtlas* available at https://github.com/cejuliano/neuron_single_cell.

scRNA-seq libraries were built using the Chromium Next GEM Single Cell 3’ Kit v3.1 (10x Genomics) at the UC Davis Genome Center with a target recovery of 10,000 cells. Final cell counts were: 11,183 for *Tg*(*tba1c:mNeonGreen*)⁽ᶜʲ¹⁻ᶠ¹⁾, and 7,988, 7,374, and 6,124 for the three *Tg*(*actin1:GFP*)⁽ʳˢ³⁻ⁱⁿ⁾ libraries. scRNA-seq libraries generated with the 10x Genomics Chromium platform were processed using Cell Ranger v4.0.0 following the manufacturer’s instructions and aligned to the *Hydra vulgaris* AEP transcriptome (Cazet et al., 2023). The average alignment rate across libraries was 59.9%, with individual alignment rates of 62.5% (*Tg(tba1c:mNeonGreen)^cj1-gt^*), 58.6% (*Tg(actin1:GFP)^rs3^* non-budding), 60.4% (budding1), and 58.1% (budding2).

Previously published scRNA-seq data from Siebert et al. (2019), mapped against the same AEP reference, were subsetted to include interstitial stem cells, neuronal progenitors, and differentiated neurons based on cluster identity. All downstream analyses were conducted using Seurat v4.0.5 (Butler et al., 2018; Hao et al., 2021; Satija et al., 2015; Stuart et al., 2019).

Individual Seurat objects were generated per library and filtered to retain cells with 300–7,000 detected genes, 500–50,000 transcripts, and <5% mitochondrial reads (Satija et al., 2015; Siebert et al., 2019). Data quality was assessed with violin plots (**Fig. S3**). For Tg(actin1:GFP)^rs3-in^ libraries, which contain a minor population of gland cells and nematocytes, clustering was performed and clusters expressing canonical markers for these cell types were identified and excluded (Siebert et al., 2019). To account for technical variation, module scores were calculated for a curated set of “stress markers” and regressed during dataset integration.

An initial dataset integration was performed using the SCTransform workflow (Hafemeister and Satija, 2019), incorporating twelve previously published whole-animal Drop-seq libraries (subsetted for interstitial stem cells, neurons, and neural progenitors), two published Drop-seq libraries of sorted neurons from the Tg(actin1:GFP)⁽ʳˢ³⁾ line, and the four newly generated 10X Genomics libraries in this study. SCTransform was first used to normalize the datasets, with regression against stress marker module scores and the percentage of mitochondrial RNA reads. Integration was performed with Seurat::SelectIntegrationFeatures (nfeatures = 3000), PrepSCTIntegration, FindIntegrationAnchors, and IntegrateData. Dimensionality reduction was carried out with Seurat::RunPCA. The number of principal components (PCs) retained was based on the point where the percent variance change between PCs dropped below 0.1%. A preliminary clustering and UMAP was generated using the Louvain approach [Seurat::*FindClusters*, (*dims.use* = 1:25), Seurat::*FindNeighbors* (*resolution* = 0.8), Seurat::*RunUMAP*, (*dims.use* = 1:25)]. One cluster expressing stress markers, mitochondrial genes, or non-neuronal lineage markers was excluded.

Following removal of the non-neuronal and high-stress cluster, datasets were reintegrated using the same SCTransform workflow. PCA was repeated, and clustering was performed with 21 PCs (resolution = 0.8). A new UMAP was generated using RunUMAP (min.dist = 0.25, spread = 0.7). Clusters were annotated based on expression of known markers for neuron subtypes, progenitors, and ISCs. Of the resulting clusters, four could not be definitively identified. Marker expression from these clusters was cross-referenced with whole-animal scRNA-seq datasets (Cazet et al., 2023; Siebert et al., 2019). One unknown cluster was enriched for stress genes and excluded. The remaining three expressed markers associated with ec1A, ec1B, and ec5 neurons and were retained. We ran Seurat::FindAllMarkers() to identify molecular markers for the 3 remaining novel clusters and validated these novel subtype markers by FISH (see below).

### ATAC-seq Library Preparation, Sequencing, and Analysis

Neuron-enriched ATAC-seq libraries were generated using a modified OMNI-ATAC protocol (Cazet et al., 2023; Corces et al., 2017; Siebert et al., 2019). FACS-isolated cells (described above) were pelleted at 1,000 g for 5 min at 4°C and resuspended in 50 µL freshly prepared, chilled resuspension buffer (10 mM Tris-HCl pH 7.4, 10 mM NaCl, 3 mM MgCl₂) supplemented with 0.1% Tween-20, 0.1% NP-40, and 0.01% digitonin. After 3 minutes on ice, lysis was quenched with 1 mL chilled RSB + 0.1% Tween-20, inverting tubes 3x to mix. Nuclei were pelleted at 500 g for 10 min at 4°C and resuspended in tagmentation mix [1x TD buffer (Illumina #20034197), 33% PBS, 0.01% digitonin, 0.1% Tween-20, 5 µL TDE1 (Illumina)]. Samples were tagmented for 30 min at 37°C in a ThermoShaker (1,000 RPM), quenched with 250 µL Qiagen PB buffer (MinElute Kit #28004), and stored at −20°C.

Tagmented DNA was purified using the Qiagen MinElute Kit, eluted in 21 µL EB buffer, and amplified with NEBNext 2X Master Mix (NEB M0541S). PCR cycle numbers were determined by qPCR (Buenrostro et al., 2015) (**Table S5**). Libraries were subjected to two rounds of size-selected (100–700 bp) using AMPure XP beads[HM1] (Beckman Coulter #A63881), quantified with Qubit dsDNA HS Assay (ThermoFisher #Q32851), and assessed with a Bioanalyzer High-Sensitivity DNA Kit (Agilent #5067-4626) at the UC Davis DNA Core. Based on concentration and trace quality, two of five Tg(tba1c:mNeonGreen)cj1-gt replicates and four of six Tg(actin1:GFP)rs3-in replicates were selected for sequencing. Libraries were pooled and sequenced using 2×150 bp reads on an Illumina HiSeq 4000 (*Tg(tba1c:mNeonGreen)cj1-gt*) or HiSeq X Ten (*Tg(actin1:GFP)rs3-in*).

A step-by-step description of the ATAC-seq analysis, including all relevant code, is provided in the markdown document *03_hydraNeuronRegulation* available at https://github.com/cejuliano/neuron_single_cell.

To analyze the ATAC-seq data collected from whole strain AEP *H. vulgaris* polyps, we first filtered the raw reads using Trimmomatic (Bolger et al., 2014) to remove stretches of low-quality base-calls and contaminating adapter sequence. The filtered reads were then aligned to the AEP assembly using Bowtie2 (Langmead and Salzberg, 2012). To remove mitochondrial reads, we also aligned the ATAC-seq data to the *H. vulgaris* mitochondrial genome (Voigt et al., 2008) and subsequently discarded any reads that aligned to the mitochondrial and nuclear genome references using Picard Tools (<u>broadinstitute.github.io/picard/</u>). We next identified and removed PCR duplicates from the aligned data using Samtools (Li et al., 2009) and Picard Tools. Peaks were called using a pipeline adapted from Cazet et al. (2023), originally based on the ENCODE ATAC-seq workflow (Landt et al., 2012). We called peaks for each ATAC-seq biological replicate using MACS2 (Zhang et al., 2008). To generate a consensus peakset of biologically reproducible ATAC-seq peaks, we first calculated irreproducible discovery rate (IDR) (Li et al., 2011) peak scores for each pairwise combination of biological replicates. We defined a reproducible peak as one that received an IDR score ≤ 0.1 for at least two pairwise comparisons between biological replicates. TSS enrichment, self-consistency, and rescue ratios were computed for quality control (Cazet et al., 2023; Siebert et al., 2019). Neuron-enriched libraries were of comparable quality to whole-animal ATAC-seq libraries (Cazet et al., 2023) (**Table S5**).

To identify differentially accessible regions, read counts across all peaks were quantified using DiffBind and annotated to the nearest transcription start site (TSS) with UROPA (Kondili et al., 2017). Differential accessibility between *Tg(tba1c:mNeonGreen)cj1-gt*, *Tg(actin1:GFP)rs3-in*, and AEP whole animal datasets (Cazet et al., 2023) was assessed using edgeR (Robinson et al., 2010). Results were saved as RData objects and summary tables, and genomic tracks were visualized using Gviz (Hahne and Ivanek, 2016). To assess whether neuron-enriched ATAC-seq libraries show greater accessibility near the TSS of neuronal genes, we first identified genes enriched (NeuroG) or depleted (nonNeuroG) in differentiated neurons using Seurat analysis of *H. vulgaris* whole-animal scRNA-seq data (Cazet et al., 2023). ATAC peaks associated with these genes were extracted from edgeR comparisons between *Tg(tba1c:mNeonGreen)cj1-gt*/whole animal AEP and *Tg(actin1:GFP)rs3-in* /whole animal AEP datasets. Log fold changes for NeuroG- and nonNeuroG-associated peaks were visualized using ggplot2 and compared using a two-sided t-test to evaluate statistical significance.

### Generation of Developmental Trajectories

A step-by-step description of the developmental trajectory analysis, including all relevant code, is provided in the markdown document *02_hydraNeuronTree* available at https://github.com/cejuliano/neuron_single_cell.

To reconstruct neuronal differentiation trajectories in Hydra, we used URD (Farrell et al., 2018) to trace the emergence of 11 neuron subtypes from interstitial stem cells (ISCs). Initial analyses revealed strong batch effects between 10x Genomics and Drop-seq datasets that impaired proper cell-to-cell transition mapping. Given that the Drop-seq data represented a minority of cells (∼5,400), we excluded it from trajectory inference and focused solely on the higher-resolution 10x scRNA-seq data. We used the SeuratToURDv3 function to generate an URD object, and manually added in the integrated data from our Seurat object (urd.obj@logupx.data <-seurat.obj@assays$integrated@data). Next, we used the URD::urdSubset function to eliminate the Drop-seq data and cell clusters that pilot analyses demonstrated lacked clear intermediate progenitors linking them back to the ISCs (ec1C, ec1D, and ec1E clusters).

Outliers, cells with poor connectivity, were removed based on k-nearest neighbor distances using URD::knnOutliers with parameters: x.max = 86, slope.r = 0.12, int.r = 79, slope.b = 1.1, and int.b = 7.75 (209 cells removed). Doublets were identified based on high co-expression of mutually exclusive gene modules (Siebert et al., 2019). NMF module scores were scaled between 0-1, and visualized on the UMAP to confirm cell-type specificity (**Fig. S6**). Doublets were called using URD::NMFDoubletsDetermineCells with parameters: module.expressed.thresh = 0.2, frac.overlap.max = 0.07, frac.overlap.diff.max = 0.15. This process flagged 1,257 potential cell doublets, which were subsequently removed.

Pseudotime trajectories were computed using a diffusion map (URD::CalcDM, with knn = 100, sigma.use = local, distance = cosine), optimized for high progenitor–differentiated cell connectivity and low noise across unrelated terminal branches. The ISC population (identified by Hy-icell1 marker G002332) was designated as the root (Siebert et al., 2019). Terminal tips were defined using Infomap-Jaccard clustering (URD::graphClustering, num.nn = 120), and selected based on late pseudotime and differential gene expression.

Pseudotime was calculated using URD::floodPseudotime (n = 50, minimum.cells.flooded = 2), and transition matrix bias was tuned with optimal.cells.forward = 0, max.cells.back = 200 to allow transitions through densely populated differentiated states. Biased random walks were performed (n.per.tip = 50,000, root.visits = 1), and visitation patterns were checked on UMAPs. In cases of multiple tips per subtype, they were merged using URD::combineTipVisitation. Final tree construction was carried out with URD::buildTree using: divergence.method = “preference”, save.all.breakpoint.info = TRUE, cells.per.pseudotime.bin = 25, bins.per.pseudotime.window = 8, p.thresh = 1e-6, and min.cells.per.segment = 10.

To examine the temporal dynamics of gene expression during differentiation, we used spline-based modeling. Genes expressed in at least 1% of cells were considered. Expression profiles across pseudotime were fit and visualized with URD::geneSmoothFit (spar = 0.875) within lineage-specific segments.

### Identification of gene modules using non-negative matrix factorization (NMF)

To identify gene co-expression modules (i.e., metagenes) in our neuronal scRNA-seq dataset, we performed non-negative matrix factorization (NMF) using the cNMF package (Kotliar et al., 2019) (Kotliar et al., 2019). We used raw read counts from all neuronal and interstitial stem cell transcriptomes across both 10x Genomics and Drop-seq libraries. Prior to factorization, data were normalized and filtered using the ‘prepare’ function to exclude genes with low variability across cells. Because the optimal number of modules (*k*) must be empirically determined, we used the ‘factorize’ function to perform independent NMF analyses for *k* values ranging from 5 to 100 in increments of 5. For each *k*, 200 independent runs were performed to assess metagene reproducibility. Results were compiled using the ‘combine’ function and evaluated for robustness and accuracy with the ‘*k*_selection_plot’ function. This initial screen suggested that the most stable solutions fell between *k* = 25 and 35. To refine our selection, we repeated the analysis for all individual k values within that range. Based on this finer-grained analysis, we selected *k* = 27 as the optimal number of modules. We then used the ‘consensus’ function to generate a final set of 27 consensus metagenes by filtering out runs that deviated most from the majority of independent solutions.

### Probe Generation for Fluorescent in Situ Hybridization

To generate labeled RNA probes, PCR products (**Table S7**) were amplified from oligo-dT-primed cDNA derived from *Hydra vulgaris* AEP. Reverse primers included T7 or SP6 promoters (Qiagen #28506). Amplicons were cloned into the Invitrogen Zero Blunt vector (#K2700-20), and correct inserts were confirmed by colony PCR and Sanger sequencing. Plasmids were amplified via miniprep (Qiagen #27106) and used as templates for a second round of PCR to generate transcription templates. Approximately 250 ng of gel-purified product was used as a transcription template for the Roche DIG RNA Labeling Kit (Sigma #11175025910) with DIG-U-11 or FITC-U-12 labeling mix (Sigma #11685619910). RNAse-free conditions were maintained using Ambion RNAse-in, and resulting RNA probes were purified with the Zymogen RNA Clean & Concentrator-25 kit (Zymo Research #R1017), diluted to 750 ng in 7.5 μL H₂O, and stored at –80°C.

### Fluorescent in Situ Hybridization

#### Fixation and Bleaching (Day 1)

Starved *Hydra vulgaris* AEP polyps were washed in *Hydra* medium (HM), relaxed in 2% urethane in HM for one minute, then fixed in 4% paraformaldehyde (PFA) in HM for 1 hour at room temperature (RT) with rocking. After fixation, samples were washed 3 × 10 min in PBT (0.1% Tween-20 in PBS) and dehydrated in a stepwise methanol series to 100% MeOH, followed by bleaching overnight at –20°C in fresh 100% MeOH.

#### Rehydration, Permeabilization, and Probe Hybridization (Day2)

Samples were rehydrated through 66% and 33% MeOH/PBT washes, followed by 3 × 5 min washes in PBT. Tissue was permeabilized with 10 μg/mL proteinase K in PBT for 5 minutes, quenched in 4 mg/mL glycine, and washed 3 × 5 min in PBT. Samples were then washed sequentially in 0.1 M triethanolamine in PBT ± acetic anhydride (3 and 6 μL/mL), refixed in 4% PFA (1 hour), washed in PBT, and equilibrated in 2× SSC. Samples were prehybridized in 50% 2× SSC/50% hybridization solution (HS: 50% formamide, 5× SSC, 1× Denhardt’s, 100 μg/mL heparin, 0.1% Tween-20, 0.1% CHAPS), first at RT then at 56°C. They were incubated for 2 hours in HS with 10 μL/mL sheared salmon sperm DNA. For sample, ∼180 ng of DIG-and/or FITC-labeled RNA was denatured at 85°C for 5–10 min, cooled to 56°C, and added to 400 μL HS. Samples were hybridized in 450 μL total probe mix at 56°C for ∼65 hours, with gentle mixing once every 24 hours.

#### Probe Washes and Antibody Staining (Day3)

Samples were washed at 56°C with decreasing HS/SSC concentrations (100%, 75%, 50%, 25%) followed by 2 × 30 min washes in 2× SSC + 0.1% CHAPS (first at 56°C, then RT). Samples were washed 4 × 10 min in MABT (100 mM maleic acid, 150 mM NaCl, 0.1% Tween-20, pH 7.5), then blocked in MABT + 1% BSA for 1 hour and in blocking buffer (80% MABT + 1% BSA / 20% sheep serum) for 2 hours at 4°C. Anti-FITC-POD (Sigma #11426346910) or Anti-DIG-POD (Sigma #11207733910) was added at 1:2000 in blocking buffer and incubated overnight at 4°C.

#### Tyramide Detection and Second Antibody Application (Day 4)

After returning to RT, excess antibody was removed with 2 × 20 min MABT-BSA and 5 × 20 min MABT washes. Samples were washed 2 × 5 min in 100 mM borate buffer + 0.1% Tween, then stained with 75 μL tyramide solution (100 mM borate buffer, 2% dextran sulfate, 0.1% Tween-20, 0.003% H₂O₂, 0.15 mg/mL 4-iodophenol, 1:100 Alexa Fluor 488 or 594 tyramide) for 25 minutes. The reaction was stopped with 4 × PBT washes and quenched in 100 mM glycine (pH 2.0) for 10 minutes, then washed 5 × 5 min in PBT. At this point, samples could either be mounted for imaging (see below) or processed for double FISH. For double labeling, samples were blocked for 2 hours at 4°C in blocking buffer (80% MABT + 1% BSA / 20% sheep serum), then incubated overnight at 4°C in 1:2000 Anti-DIG-POD (Sigma #11207733910) in blocking buffer, protected from light. <u>Second Tyramide</u> <u>Detection and Mounting (Day 5)</u>: Following antibody incubation, samples were washed and processed for the second round of tyramide detection as described above, using the alternate fluor. After staining, samples were again quenched in 100 mM glycine (pH 2.0) and washed 5 × 5 minutes in PBT. To prepare for imaging, tissues were stained with 1:1000 Hoechst for 30 minutes, then dehydrated through a graded series of glycerol (30%, 50%, 80% in PBT) for at least 1 hour in each step. Specimens were mounted in 80% glycerol and stored at 4°C until imaging. Confocal images were acquired using the Zeiss 980 LSM with Airyscan 2 confocal microscope. For whole animal images, max projections were stitched together using ImageJ.

#### In Situ Hybridization Chain Reaction

The ec1E cluster could not be identified using traditional FISH and instead required detection via In Situ Hybridization Chain Reaction (HCR) (Choi et al., 2018). <u>Probe Design and Synthesis</u>: probes were designed using the in situ_probe_generator tool (https://github.com/guttalab/in_situ_probe_generator) (Kuehn et al., 2022). Probes were designed to target transcripts of interest and synthesized by IDT. <u>Sample Preparation, Fixation,</u> <u>and Permeabilization</u>: *Hydra* were washed twice in sterile *Hydra* Medium (HM) and relaxed in 2% urethane in HM for 1 minute. Samples were fixed in 4% paraformaldehyde (PFA) in HM for 1 hour with gentle rocking. Fixed samples were washed three times for 10 minutes each in phosphate-buffered saline with Tween (PBT), followed by gradual dehydration through 5-minute washes in increasing concentrations of methanol (MeOH) in PBT: 33%, 66%, and 100%.

Samples were then washed twice for 30 minutes in 100% MeOH and stored in fresh 100% MeOH at −20°C overnight. <u>Probe Hybridization</u>: Samples were rehydrated by sequential 5-minute washes in 66% MeOH in water and 33% MeOH in PBT, followed by three 5-minute washes in PBT. Samples were then incubated in 1ml of 50% probe hybridization buffer in PBT for 5 minutes, followed by incubation in 300µl of 100% probe hybridization buffer for 1 hour at 37°C. Probes (1–2 pmol each) were diluted into pre-warmed hybridization buffer (250µl total volume), and samples were incubated in this hybridization mix overnight at 37°C. <u>Probe</u> <u>Detection and Signal Amplification</u>: Following hybridization, samples were washed four times for 15 minutes each with 500µl of probe wash buffer pre-warmed to 37°C. Samples were then washed three times for 5 minutes each in 5x SSC with 0.1% Tween-20 (SSCT) at room temperature, followed by a 5-minute wash in PBT. Samples were equilibrated in 300µl of amplification buffer for 30 minutes. Meanwhile, hairpin amplifiers were prepared by heating each pair of hairpins separately to 95°C for 90 seconds, followed by cooling to room temperature in the dark for 30 minutes. The hairpins (15 pmol each) were then combined in amplification buffer (250µl total volume), and samples were incubated in the hairpin mix overnight in the dark. <u>Post-Amplification Washes and Mounting</u>: Following amplification, samples were washed twice for 5 minutes and twice for 30 minutes in 5x SSCT. DAPI was optionally added during the first 30-minute wash. Samples were stored in 5x SSCT at 4°C until imaging. Prior to imaging, samples were washed twice for 5 minutes in PBS and mounted in ProLong Glass mounting medium. Confocal images were acquired using the Zeiss 980 LSM with Airyscan 2 confocal microscope.

#### Immunofluorescence

*Hydra* polyps were anesthetized in 2% urethane in *Hydra* medium for 4–5 minutes, then fixed in 4% paraformaldehyde (PFA) in 1× *Hydra* PBS (120 mM NaCl, 40 mM K2HPO4, 12 mM NaH2PO4; pH 7.2-7.4) for 1 hour at room temperature. Following fixation, animals were washed three times with 1× *Hydra* PBS for 10 minutes each on a shaker. Fixed animals were then cut as needed and permeabilized in 0.5% Triton X-100 in 1× *Hydra* PBS for 15 minutes in the dark. Blocking solution was prepared by dissolving 0.05 g BSA and 50 µL 10% Triton X-100 in 4950 µL 1× *Hydra* PBS, and animals were incubated in this solution for 20 minutes in the dark. Primary antibodies were diluted in blocking solution and applied at the following concentrations: pan-neural antibody (1:1000, raised in rabbit) (Keramidioti et al., 2024), anti-GFP (1:500, raised in chicken, Aves Labs, GFP-1020), anti-mNeonGreen (1:50, raised in mouse, Chromotek, 32f6), anti-GLWa (1:10, raised in mouse) (Koizumi et al., 2015), and anti-RFa (1:50, raised in mouse) (Koizumi et al., 2015). 100 µL of the appropriate antibody dilution was added per well (one animal per well), and samples were incubated overnight at 4 °C. The following day, animals were washed three times in 1× *Hydra* PBS for 10 minutes each on a shaker in the dark.

Secondary antibodies were diluted in blocking solution and included Alexa Fluor 488 donkey anti-chicken (1:400, Dianova, 703-546-155), Cy3 donkey anti-rabbit (1:400, Dianova, 711-165-152), Alexa Fluor 488 donkey anti-mouse (1:300, Dianova, 715-545-150), Cy3 donkey anti-mouse (1:200, Dianova, 715-166-151), and Alexa Fluor 647 donkey anti-mouse (1:300, Dianova, 715-605-150). 100 µL of secondary antibody solution was added per well and samples were incubated for 2 hours at room temperature in the dark, followed by three 10-minute washes in 1× *Hydra* PBS. DAPI (Sigma) was used to stain nuclei at a dilution of 1:1000 in 1× *Hydra* PBS, incubating 100 µL per well for 7 minutes. After a final 5-minute wash in 1× *Hydra* PBS, the stained specimens were mounted on microscope slides using VECTASHIELD HardSet Antifade Mounting Medium to preserve fluorescence. Bee wax was applied around the edges of the coverslip to prevent compression of the specimens, and the coverslip was sealed with transparent nail polish. Preparations were stored at 4°C until imaging. Fluorescent confocal microscopy was performed on a Leica SP5 scanning confocal microscope (Leica Microsystems, Heidelberg, Germany). Z-shift correction and RGB stack generation were performed using the StackGroom plugin in ImageJ with the “3channels” command. The resulting RGB stacks were further processed in ImageJ, and all images were generated as maximum intensity projections using the “Image > Stack > Z Project” function.

## Acknowledgments

We would like to thank Rob Steele for providing the *Tg(actin1:GFP)*^rs3-in^ line, which was foundational to this study; the Juliano lab, Farrell lab, and David lab for invaluable discussions and feedback; Bruce Draper, Rob Steele, and Stefan Siebert for manuscript review and feedback; Clara Stuligross for analysis help and moral support; Shun Hamada and Osam Koizumi for providing GLWa and RFa antibodies; Yohana Berhe for help with confocal images used in Fig. S2; Diana Burkart-Waco, Siranoosh Ashtari, and Emily Kumimoto from the DNA Technologies and Expression Analysis Core at the UC Davis Genome Center (supported by NIH Shared Instrumentation Grant 1S10OD010786-01) for technical advice and assistance with library preparation and sequencing; Bridget McLaughlin and Jonathan Van Dyke from the UC Davis Flow Cytometry Shared Resource Laboratory for technical advice and assistance with FACS [(supported by NCI P30 CA093373 (Comprehensive Cancer Center), S10 OD018223 (Astrios Cell Sorter), S10 RR 026825 (Fortessa Cytometer), and NIH NCRR C06-RR12088 (Tupper Hall room 3425 sorting suite renovations)].

## Data and resource availability

The raw sequencing data in this study have been submitted to the NCBI BioProject database (https://www.ncbi.nlm.nih.gov/bioproject/) under accession number PRJNA1261600. Step-by-step descriptions of all computational analyses conducted as part of this study, including all relevant code, formatted both as markdown and HTML documents, are available at GitHub (https://github.com/cejuliano/neuron_single_cell).

