## Supplemental Figures for "A Molecular, Spatial, and Regulatory Atlas of the *Hydra vulgaris* Nervous System"

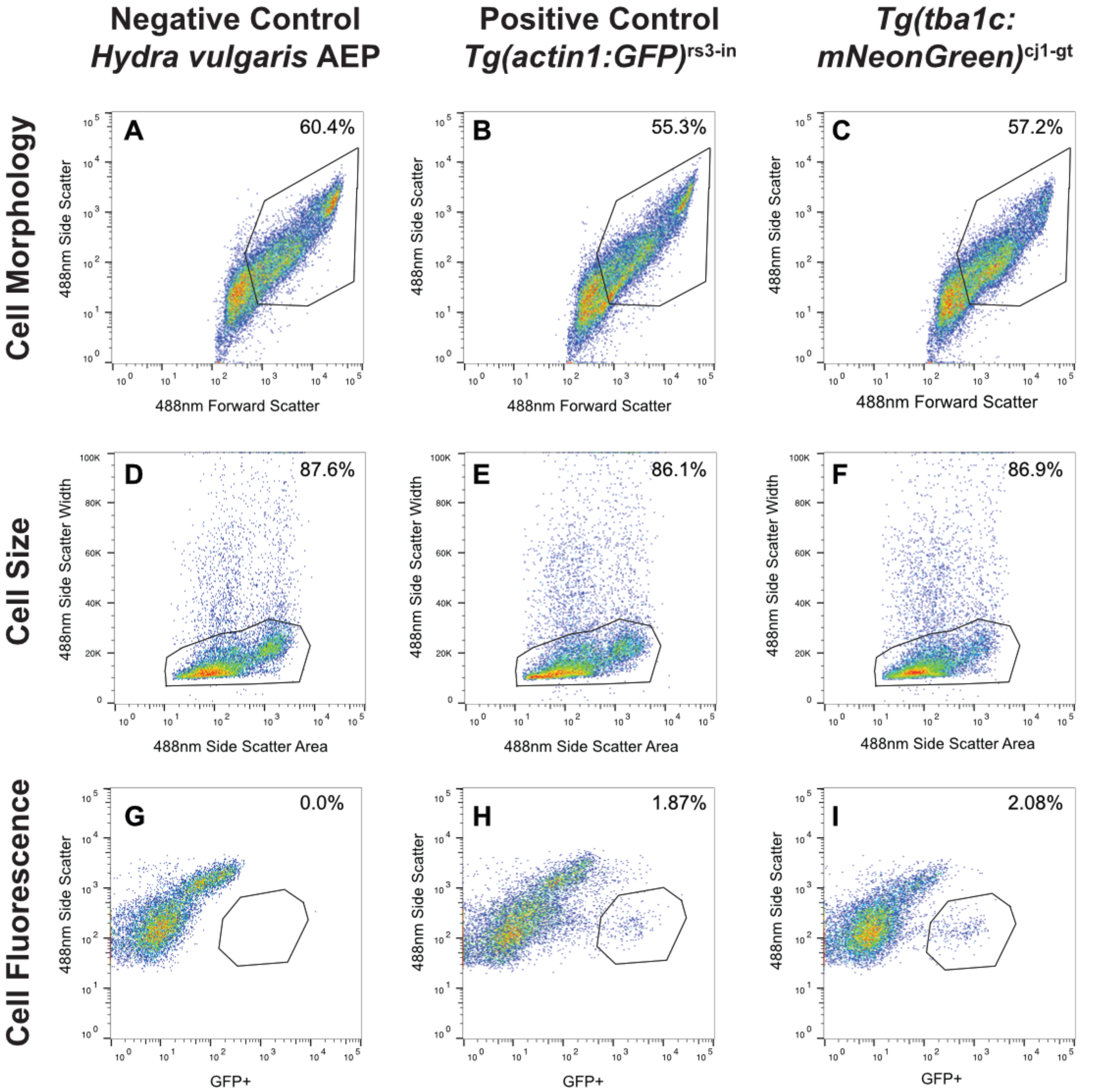

Negative Control
Hydra vulgaris AEP
Positive Control
Tg(actin1:GFP)rs3-in
Tg(tba1c:
mNeonGreen)cj1-gt
Cell Morphology
Cell Size
Cell Fluorescence
A
60.4%
B
55.3%
C
57.2%
D
87.6%
E
86.1%
F
86.9%
G
0.0%
H
1.87%
I
2.08%
488nm Side Scatter
488nm Forward Scatter
488nm Side Scatter Width
488nm Side Scatter Area
GFP+

**Figure S1. Gating for Fluorescent Activated Cell Sorting (FACS) of transgenic *H. vulgaris* cells.** This figure illustrates the gating specifications used during FACS collection of transgenic *Tg(tba1c:mNeonGreen)$^{cj1\text{-}gt}$ Hydra* cells for scRNA-seq and ATAC-seq. *Tg(actin1:GFP)$^{rs3\text{-}in}$* cells were collected using established gating parameters from Siebert et al. (2019) and served as a positive control to guide gating for this study. Gating steps included: (**A-C**) selection based on cell morphology, (**D-F**) exclusion of doublets based on cell size, and (**G-I**) selection of GFP/mNeonGreen-expressing cells using fluorescence intensity. Representative gating plots are shown for the negative FACS control *H. vulgaris* AEP (**A, D, G**), the positive FACS control *Tg(actin1:GFP)$^{rs3\text{-}in}$* (**B, E, H**) (Keramidioti et al., 2023), and *Tg(tba1c:mNeonGreen)$^{cj1\text{-}gt}$*. (**C, F, I**). The percentage of total cells within each gate is indicated in the top-right corner of each panel.

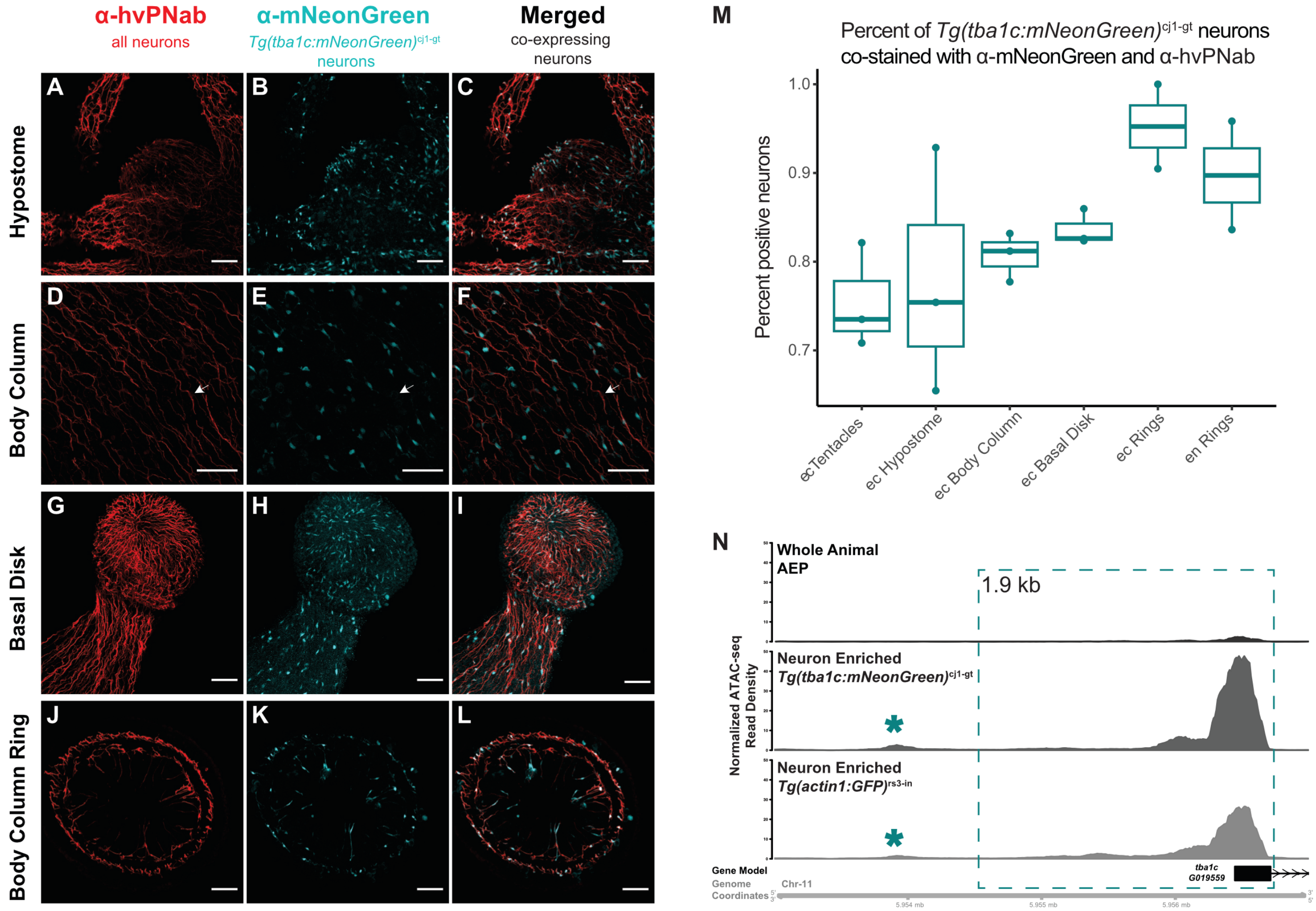
α-hvPNab
all neurons
α-mNeonGreen
Tg(tba1c:mNeonGreen)cj1-gt neurons
Merged
co-expressing neurons
A
B
C
D
E
F
G
H
I
J
K
L
Hypostome
Body Column
Basal Disk
Body Column Ring
M
Percent of Tg(tba1c:mNeonGreen)cj1-gt neurons co-stained with α-mNeonGreen and α-hvPNab
Percent positive neurons
1.0
0.9
0.8
0.7
ecTentacles
ec Hypostome
ec Body Column
ec Basal Disk
ec Rings
en Rings
N
Whole Animal AEP
1.9 kb
Neuron Enriched Tg(tba1c:mNeonGreen)cj1-gt
Neuron Enriched Tg(actin1:GFP)rs3-in
Normalized ATAC-seq Read Density
Gene Model
Genome Coordinates
Chr-11
tba1c G019559
5.954 mb
5.955 mb
5.956 mb

**Figure S2. Tg(*tba1c*:mNeonGreen)$^{cj1\text{-}gt}$ line displays mosaic transgene expression in neurons despite germline transmission.** (**A-L**) Confocal images of *Tg(tba1c:mNeonGreen)*$^{cj1\text{-}gt}$ *Hydra* immunostained with PNab (red), labeling all neurons (Keramidioti et al., 2024), and NeonGreen (cyan) fluorescent neurons, marking transgenic neurons. Representative images show different regions along the body column: (**A-C**) hypostome and tentacles, (**D-F**) ectodermal body column, (**G-I**) basal disk and (**J-L**) Cross-sectional view of excised tissue from the gastric region showing the endodermal epithelium. All transgenic neurons exhibit PNab staining, but some PNab-positive neurons lack mNeonGreen fluorescence (e.g., white arrow). Scale bar: 50 µm. **(M)** Quantification of confocal images calculating the percentage of neurons co-labeled with α-mNeonGreen and PNab across different body regions. Data were collected from n ≥3 animals per region, with 1-4 stacks analyzed per section. ec, ectodermal; en, endodermal **(N)** Chromatin accessibility at the *tba1c* locus. The Tg(*tba1c*:mNeonGreen)$^{cj1\text{-}gt}$ line was constructed using a 1,901 bp region upstream of the *tba1c* transcription start site (TSS), identified as a regulatory element from whole-animal ATAC-seq data (Siebert et al., 2019; dashed box). Neuron-enriched ATAC-seq data (**Fig. 7**) identified an additional possible regulatory peak ~3,000 bp upstream of the TSS (asterisks), that was not included in the transgenic line.

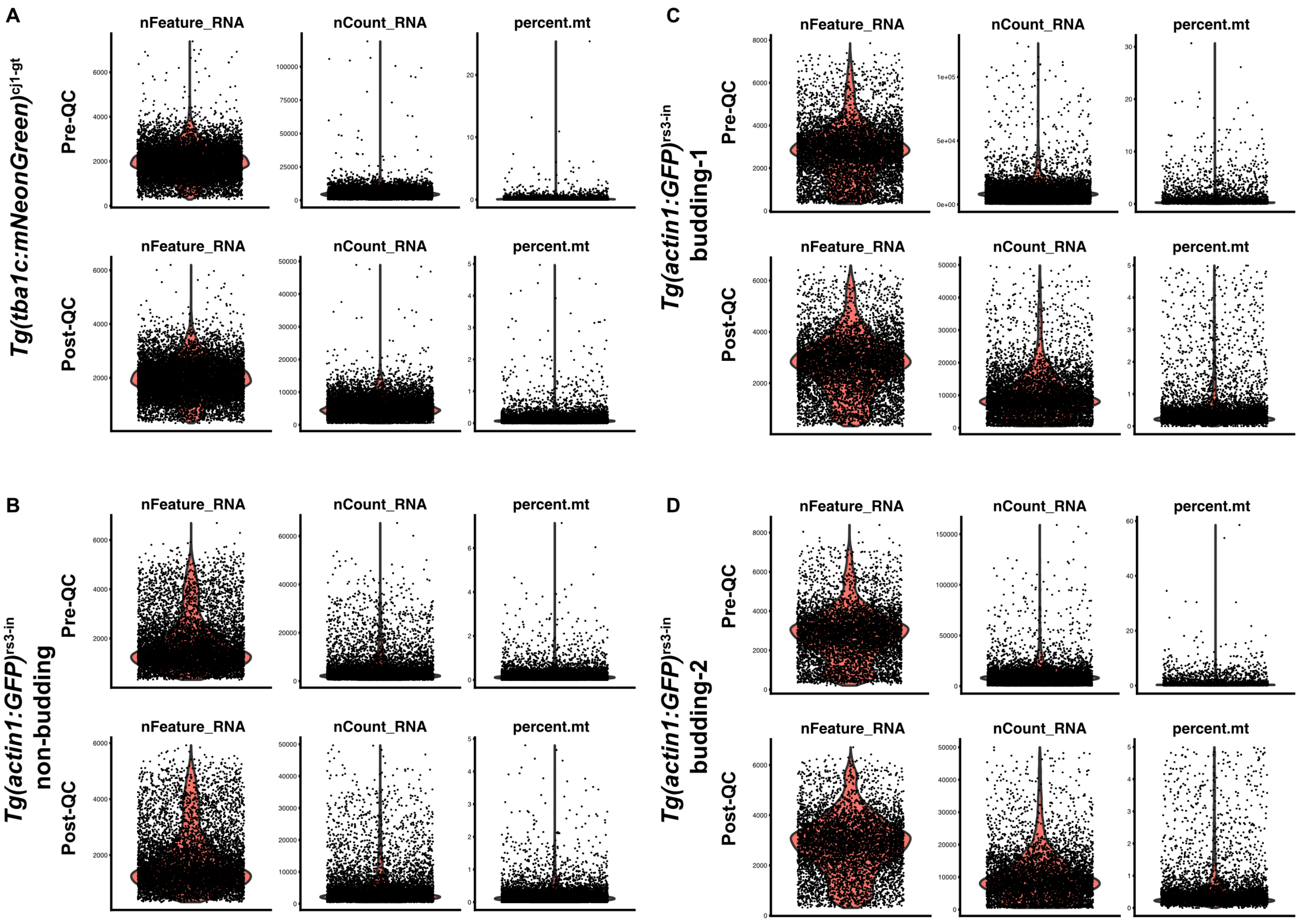

A
C
B
D
Tg(tba1c:mNeonGreen)cj1-gt
Tg(actin1:GFP)rs3-in budding-1
Tg(actin1:GFP)rs3-in non-budding
Tg(actin1:GFP)rs3-in budding-2
Pre-QC
Post-QC
nFeature_RNA
nCount_RNA
percent.mt

**Figure S3. Quality control (QC) plots for single cell RNA-seq libraries.** QC plots are shown before and after filtering (pre-QC vs. post-QC) for neuron-enriched single-cell RNA-seq (scRNA-seq) libraries generated from transgenic *Hydra* lines: (**A**) *Tg(tba1c:mNeonGreen)*$^{cj1\text{-}gt}$, (**B**) non-budding Tg(*actin1*:GFP)$^{rs3\text{-}in}$, (**C**) budding *Tg(actin1:GFP)*$^{rs3\text{-}in}$ library 1, (**D**) budding *Tg(actin1:GFP)*$^{rs3\text{-}in}$ library 2. Each dot represents a single-cell transcriptome, with corresponding violin plots displaying the distribution of the following metrics: *nFeature_RNA* refers to the number of genes detected per cell, *nCount_RNA* refers to the number of transcripts detected per cells, and *percent.mt* refers to the percentage of mitochondrial transcripts detected per cell. Cells selected for downstream analysis met the following criteria: (1) 300 - 7,000 uniquely expressed genes (*nFeature_RNA*), (2) 500 - 50,000 transcripts (*nCount_RNA*), and (3) less than 5% mitochondrial reads (*percent.mt*).

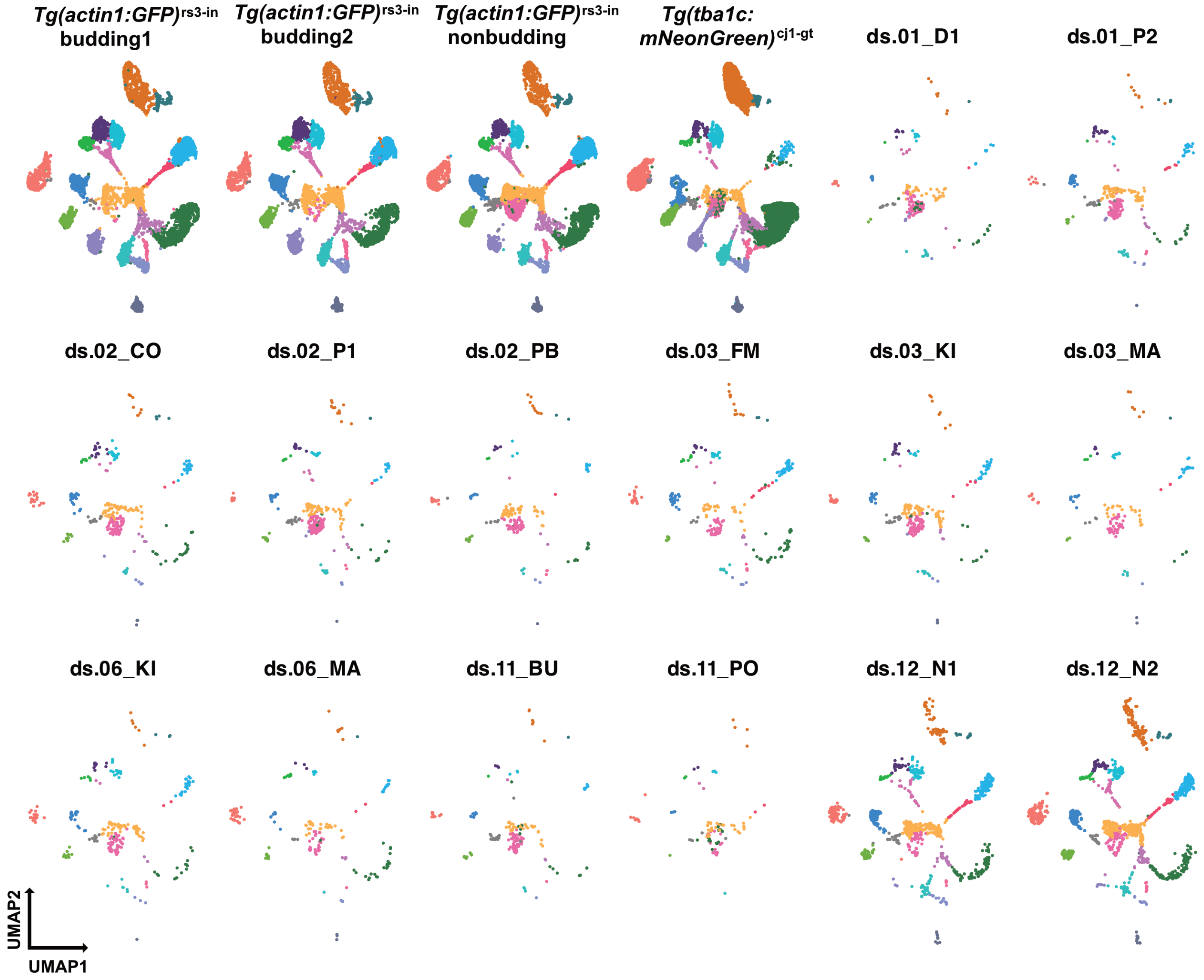

Tg(actin1:GFP)rs3-in budding1
Tg(actin1:GFP)rs3-in budding2
Tg(actin1:GFP)rs3-in nonbudding
Tg(tba1c: mNeonGreen)cj1-gt
ds.01_D1
ds.01_P2
ds.02_CO
ds.02_P1
ds.02_PB
ds.03_FM
ds.03_KI
ds.03_MA
ds.06_KI
ds.06_MA
ds.11_BU
ds.11_PO
ds.12_N1
ds.12_N2
UMAP2
UMAP1

**Figure S4. UMAP projections of individual library contributions to the combined dataset.** Detailed descriptions of the libraries are provided in Table S2.

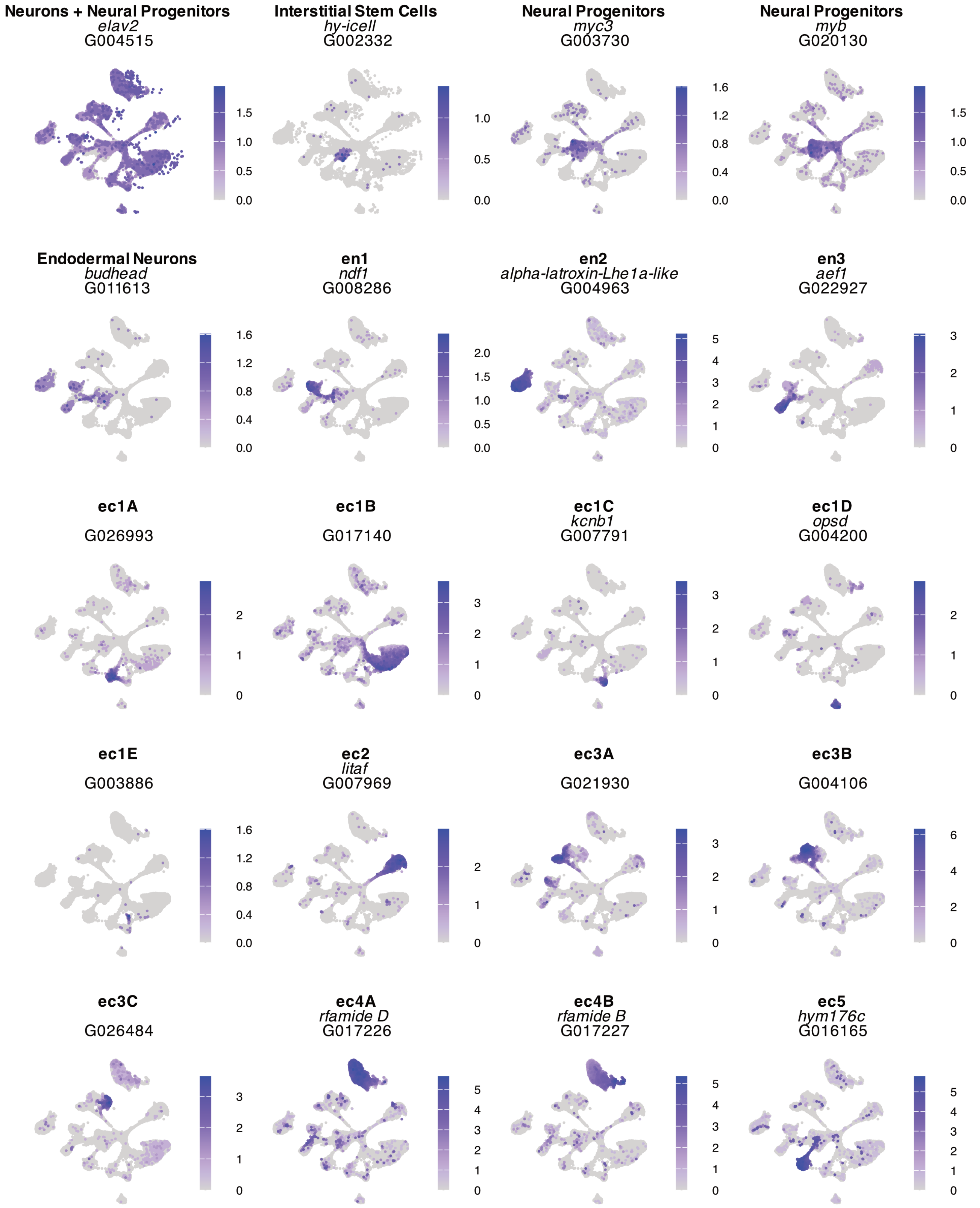
Neurons + Neural Progenitors
elav2
G004515
0.0 0.5 1.0 1.5
Interstitial Stem Cells
hy-icell
G002332
0.0 0.5 1.0
Neural Progenitors
myc3
G003730
0.0 0.4 0.8 1.2 1.6
Neural Progenitors
myb
G020130
0.0 0.5 1.0 1.5
Endodermal Neurons
budhead
G011613
0.0 0.4 0.8 1.2 1.6
en1
ndf1
G008286
0.0 0.5 1.0 1.5 2.0
en2
alpha-latroxin-Lhe1a-like
G004963
0 1 2 3 4 5
en3
aef1
G022927
0 1 2 3
ec1A
G026993
0 1 2
ec1B
G017140
0 1 2 3
ec1C
kcnb1
G007791
0 1 2 3
ec1D
opsd
G004200
0 1 2
ec1E
G003886
0.0 0.4 0.8 1.2 1.6
ec2
litaf
G007969
0 1 2
ec3A
G021930
0 1 2 3
ec3B
G004106
0 2 4 6
ec3C
G026484
0 1 2 3
ec4A
rfamide D
G017226
0 1 2 3 4 5
ec4B
rfamide B
G017227
0 1 2 3 4 5
ec5
hym176c
G016165
0 1 2 3 4 5

**Figure S5. Cell-type markers used for cluster annotations and/or FISH validation.** UMAP plots show the expression of marker genes used to annotate single-cell clusters. Markers for interstitial stem cells, progenitor cells, and endodermal neurons were identified in Siebert et al., 2019. Markers for ectodermal neurons correspond to genes used in the FISH experiments shown in Fig. 2. The title of each plot indicates the annotated cell state (in bold), with the corresponding gene name (if applicable) and the gene ID from the *Hydra vulgaris* AEP genome shown below (Cazet et al., 2023).

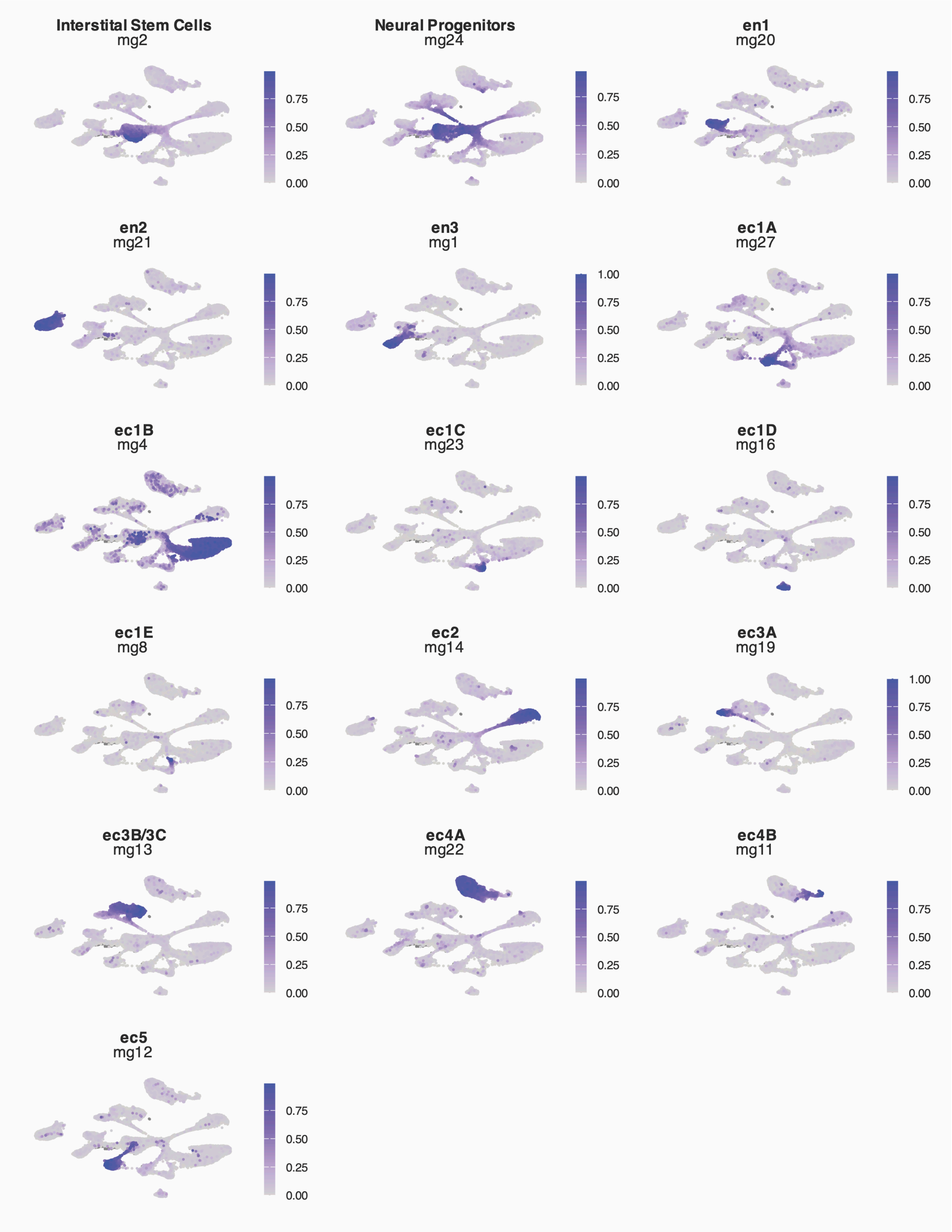
Interstital Stem Cells
mg2
0.75
0.50
0.25
0.00
Neural Progenitors
mg24
0.75
0.50
0.25
0.00
en1
mg20
0.75
0.50
0.25
0.00
en2
mg21
0.75
0.50
0.25
0.00
en3
mg1
1.00
0.75
0.50
0.25
0.00
ec1A
mg27
0.75
0.50
0.25
0.00
ec1B
mg4
0.75
0.50
0.25
0.00
ec1C
mg23
0.75
0.50
0.25
0.00
ec1D
mg16
0.75
0.50
0.25
0.00
ec1E
mg8
0.75
0.50
0.25
0.00
ec2
mg14
0.75
0.50
0.25
0.00
ec3A
mg19
1.00
0.75
0.50
0.25
0.00
ec3B/3C
mg13
0.75
0.50
0.25
0.00
ec4A
mg22
0.75
0.50
0.25
0.00
ec4B
mg11
0.75
0.50
0.25
0.00
ec5
mg12
0.75
0.50
0.25
0.00

**Figure S6. Selected metagenes identified through NMF analysis.** Metagenes represent groups of co-expressed genes identified using non-negative matrix factorization (NMF) (Kotliar et al., 2019). Instances of inappropriate metagene co-expression were used in URD analysis to identify cells likely to be doublets.

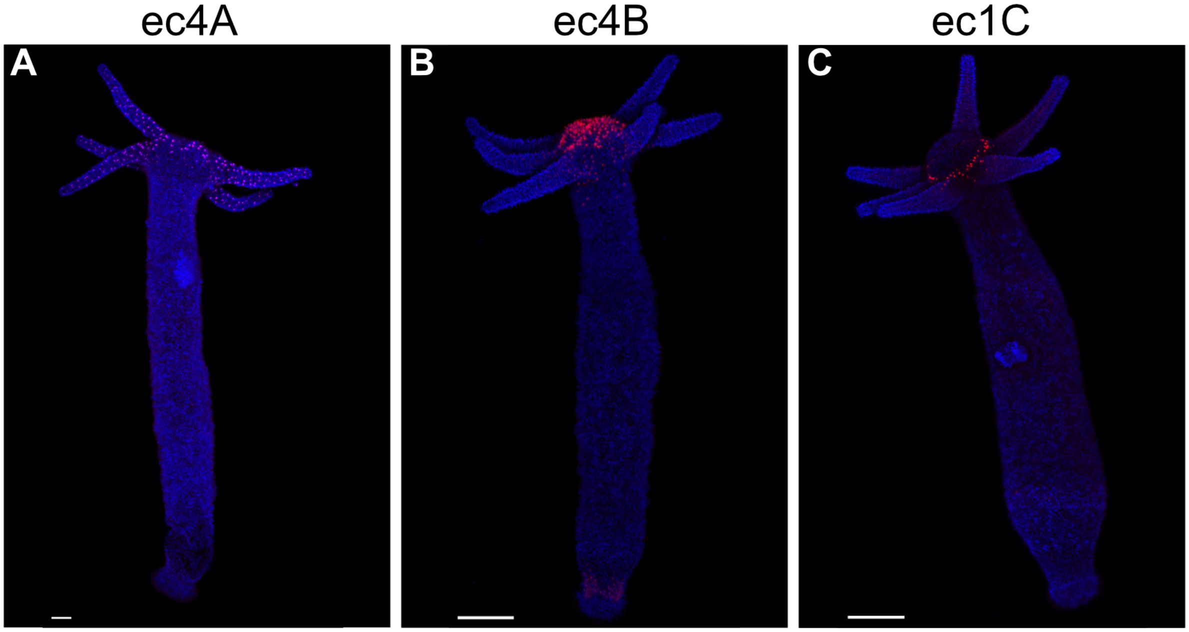
ec4A
ec4B
ec1C
A
B
C

**Figure S7. Whole animal FISH images for ec4A, ec4B, and ec1C neuron subtypes.** See Fig. 2 for zoomed in views of oral regions.

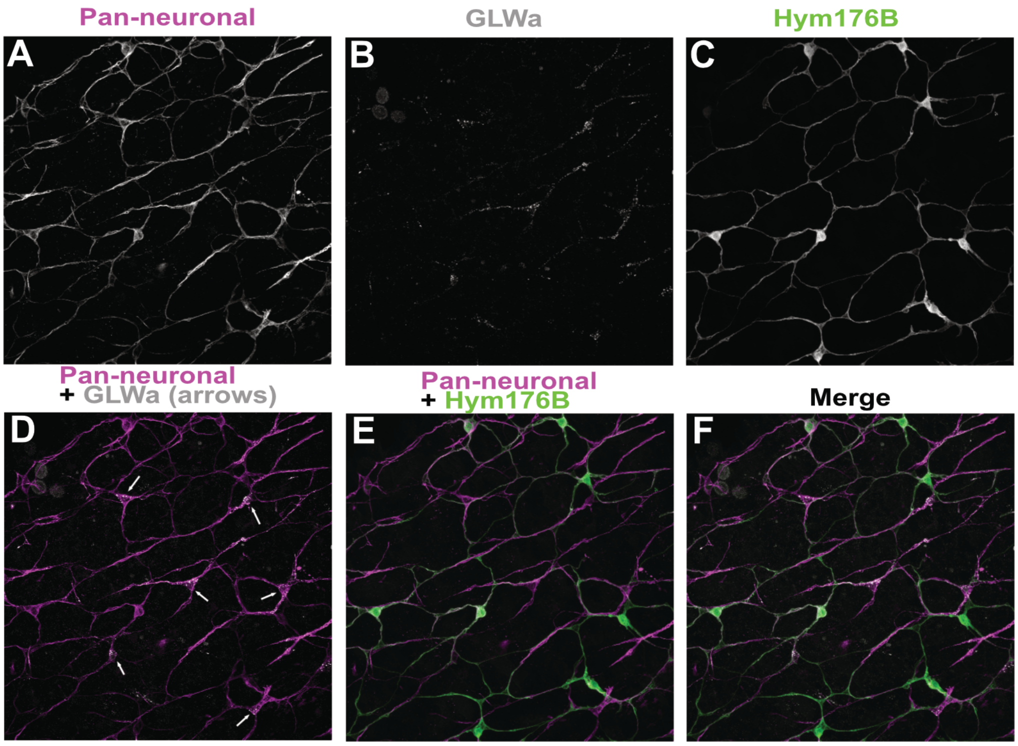
Pan-neuronal
GLWa
Hym176B
A
B
C
Pan-neuronal
+ GLWa (arrows)
Pan-neuronal
+ Hym176B
Merge
D
E
F

**Figure S8. ec1 and ec3 neuron types are the only subtypes present in the body column ectodermal nerve net.** Triple immunofluorescence was performed using antibodies against PNab (magenta, pan-neuronal marker), GLWa (white, labels ec3 neurons), and GFP (green) in the *Hym17B-GFP* reporter line (GFP labels ec1 neurons). **(A–C)** Individual channels showing PNab (A), GLWa (B), and GFP (C) staining. **(D)** Overlay of PNab and GLWa highlights GLWa-positive ec3 neurons (arrows). (E) Overlay of PNab and GFP shows GFP-positive ec1 neurons. (F) Merged image of all three channels confirms that only ec1 and ec3 neuron types are located in the body column.

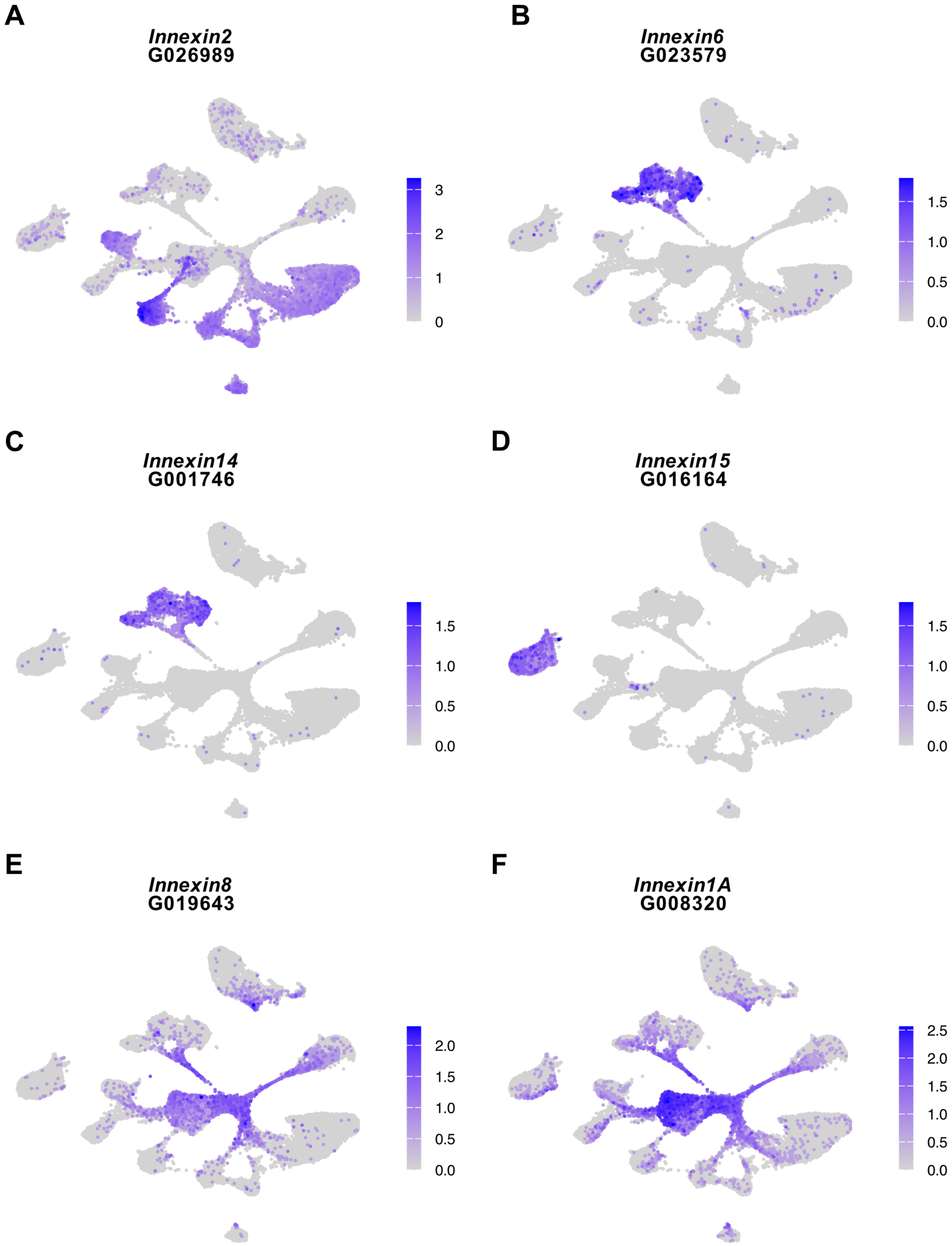
A
Innexin2
G026989
3
2
1
0
B
Innexin6
G023579
1.5
1.0
0.5
0.0
C
Innexin14
G001746
1.5
1.0
0.5
0.0
D
Innexin15
G016164
1.5
1.0
0.5
0.0
E
Innexin8
G019643
2.0
1.5
1.0
0.5
0.0
F
Innexin1A
G008320
2.5
2.0
1.5
1.0
0.5
0.0

**Figure S9. Unique innexin gene expression patterns in neuron subtypes suggest roles in circuit-specific communication.** UMAPs showing the expression of innexin genes across *H. vulgaris* neuron clusters.

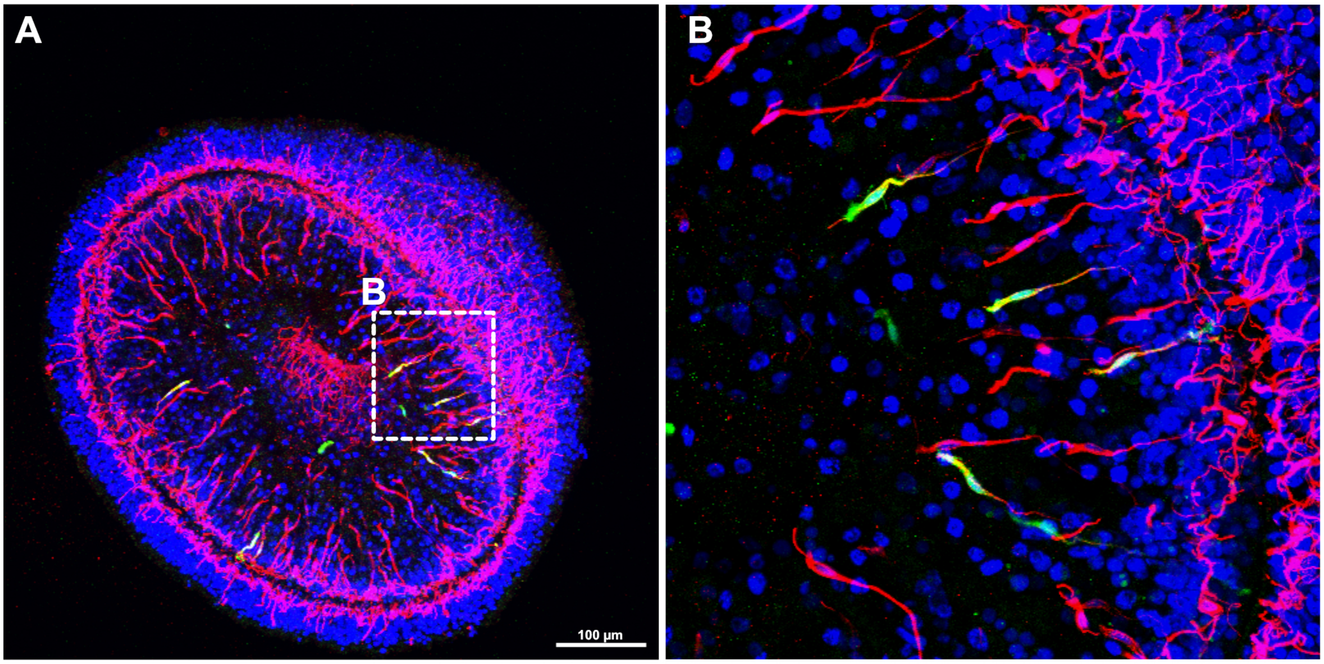
A
B
100 µm
B

**Figure S10. en3 neurons exhibit sensory neurons morphology.** A transgenic *H. vulgaris* line was generated to express mNeonGreen under the control of the en3-specific promoter (*G022927*), enabling specific labeling of en3 neurons. **(A)** Cross-sectional view of the transgenic animal showing mNeonGreen-positive cells co-stained with PNab (Keramidioti et al., 2024). **(B)** Magnified view of the boxed region in panel A. en3 neurons display morphological features characteristic of endodermal sensory neurons, including a short apical sensory cilium projecting into the gastric cavity and an elongated basal neurite extending toward ganglion neurons. Image in Panel A was taken with a 20x objective, B is the same sample taken with a 63x objective.

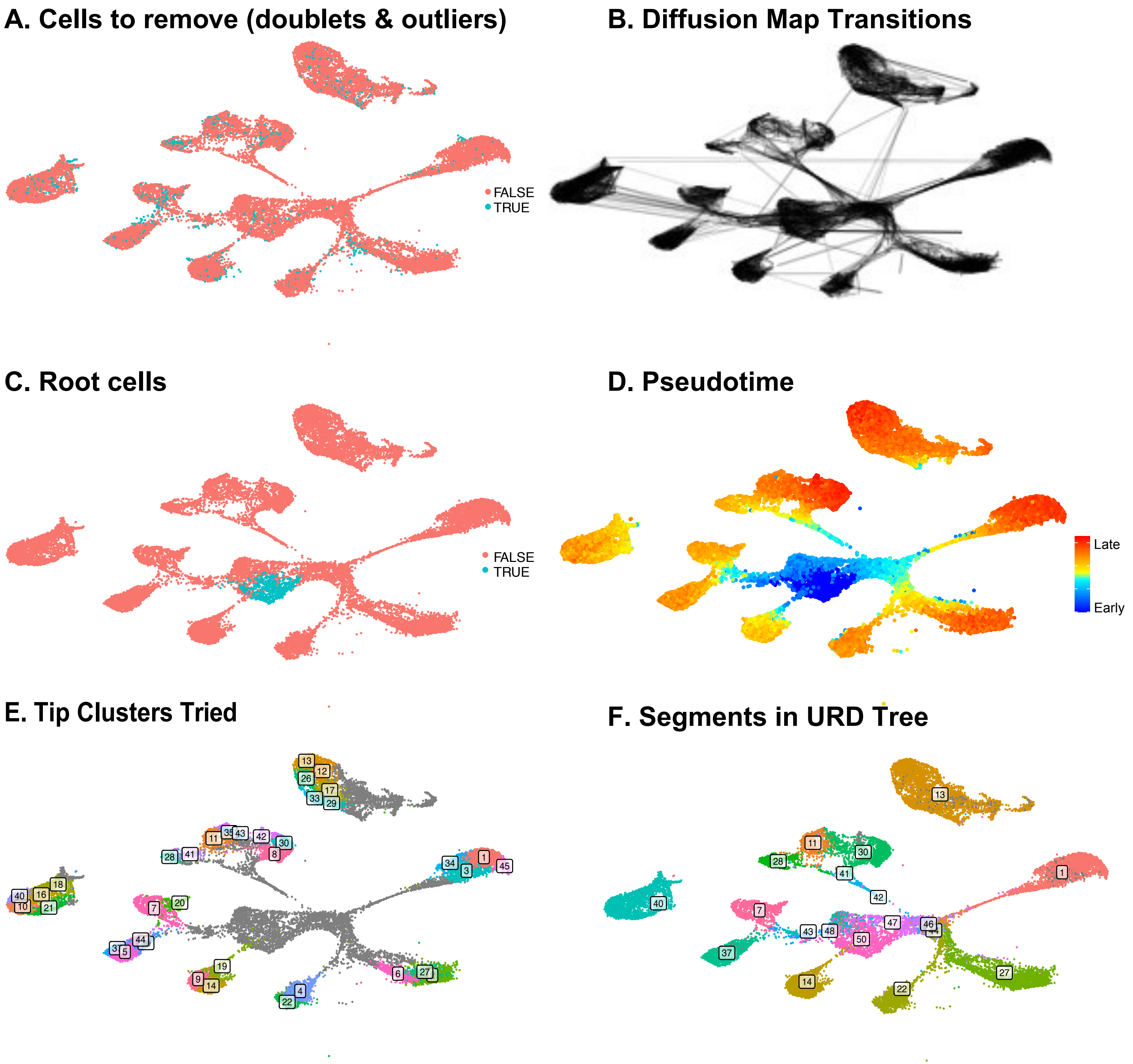
A. Cells to remove (doublets & outliers)
FALSE
TRUE
B. Diffusion Map Transitions
C. Root cells
FALSE
TRUE
D. Pseudotime
Late
Early
E. Tip Clusters Tried
F. Segments in URD Tree

**Figure S11. Trajectory Reconstruction Intermediate Steps.** Intermediate steps in URD trajectory reconstruction are represented here on the UMAP projection from Figure 1. (**A**) Cells that were identified as doublets or outliers (TRUE) were removed from consideration during trajectory reconstruction. (**B**) Diffusion map parameters were used that preserved cell-cell connections (black lines) between cells within the same cluster and a limited number of additional clusters. (**C**) Cells used as the starting point (TRUE) for trajectory reconstruction. (**D**) URD-assigned pseudotime of cells. (**E**) Cells used as end points for differentiated neuronal populations during trajectory reconstruction. (**F**) URD-assigned segment identities for cells, numbered according to the tree representation in Figure 5A.

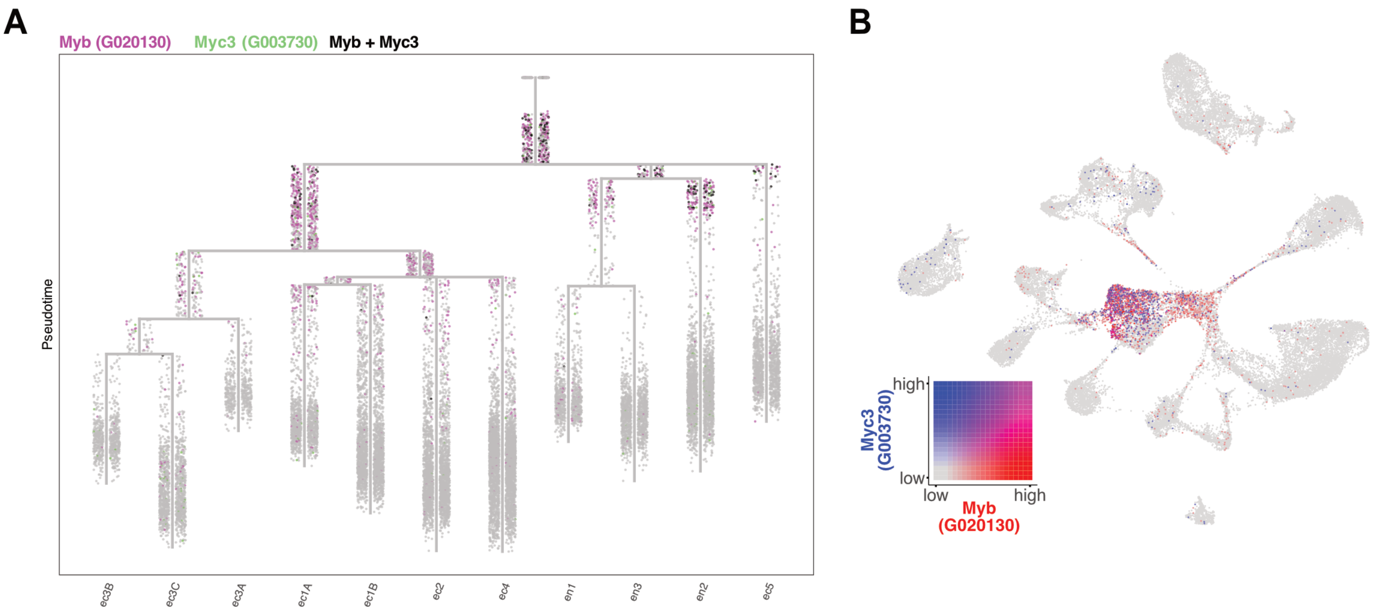

A
Myb (G020130)
Myc3 (G003730)
Myb + Myc3
Pseudotime
ec3B
ec3C
ec3A
ec1A
ec1B
ec2
ec4
en1
en3
en2
ec5
B
high
low
low
high
Myc3 (G003730)
Myb (G020130)

**Figure S12. *Myc3* and *Myb* mark early neuronal progenitor populations. (A)** Co-expression of *Myc3* (green) and *Myb* (magenta) visualized on the URD trajectory tree, with overlapping expression shown in black. This pattern highlights two distinct neuronal progenitor populations that branch early from interstitial stem cells (ISCs). **(B)** Co-expression of *Myc3* (blue) and *Myb* (red) visualized on a UMAP. These transcription factors are broadly and transiently expressed at the onset of neurogenesis and are proposed to define early neural lineage commitment.

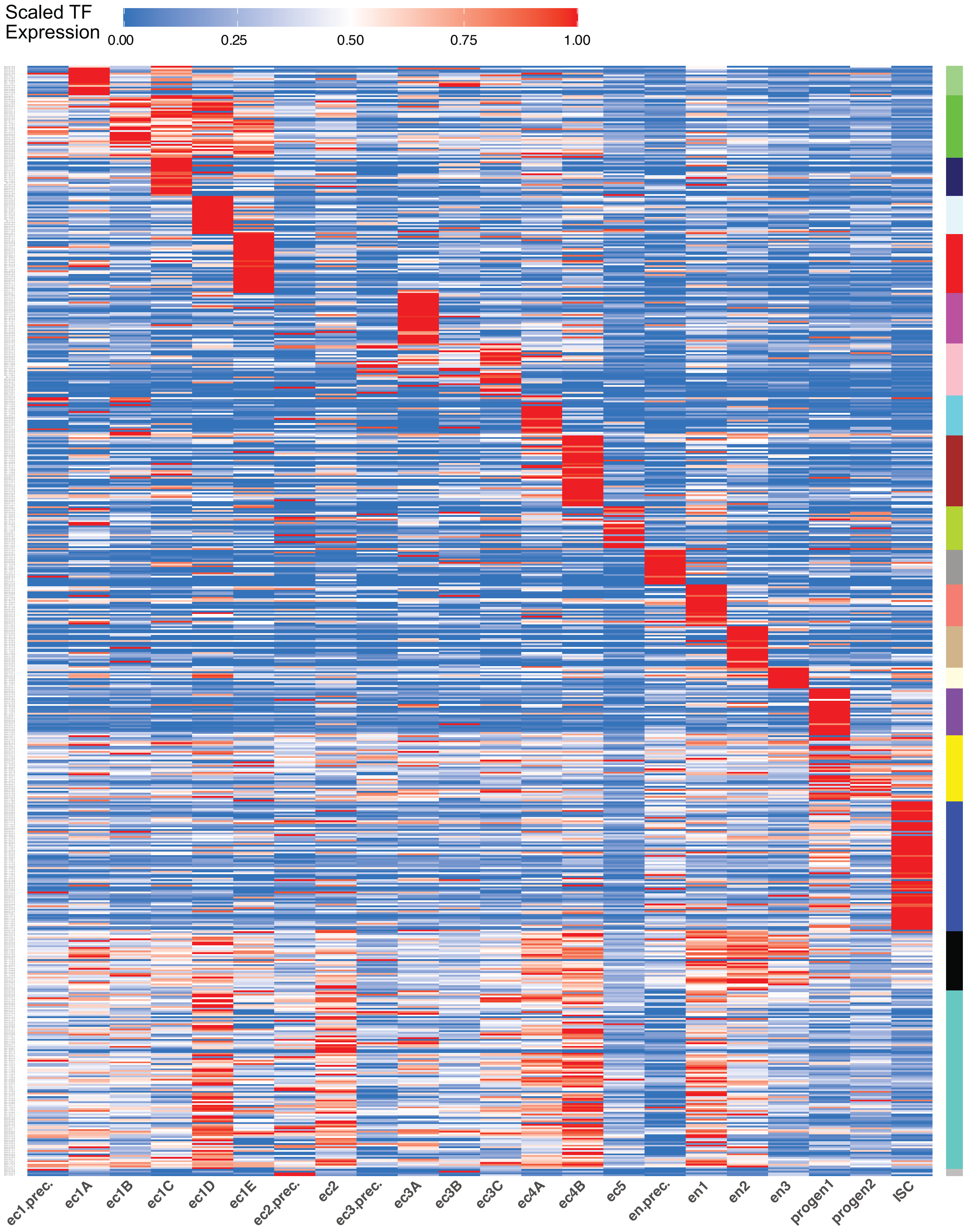

Scaled TF
Expression
0.00
0.25
0.50
0.75
1.00
ec1.prec.
ec1A
ec1B
ec1C
ec1D
ec1E
ec2.prec.
ec2
ec3.prec.
ec3A
ec3B
ec3C
ec4A
ec4B
ec5
en.prec.
en1
en2
en3
progen1
progen2
ISC

**Figure S13. Heatmap of putative transcription factors (TFs) expressed in *Hydra vulgaris* neurons.**

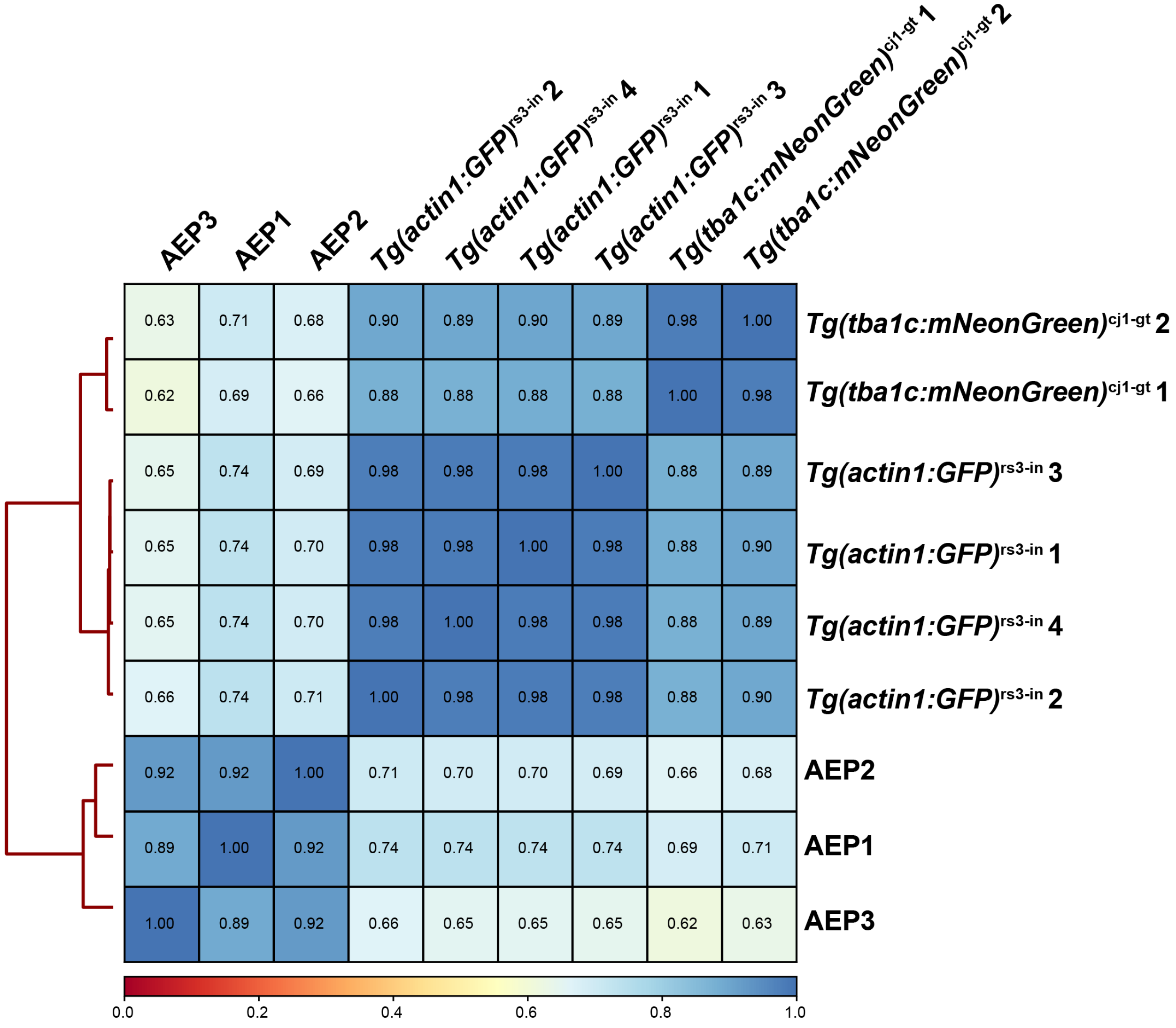

AEP3
AEP1
AEP2
Tg(actin1:GFP)rs3-in 2
Tg(actin1:GFP)rs3-in 4
Tg(actin1:GFP)rs3-in 1
Tg(actin1:GFP)rs3-in 3
Tg(tba1c:mNeonGreen)cj1-gt 1
Tg(tba1c:mNeonGreen)cj1-gt 2
0.63 0.71 0.68 0.90 0.89 0.90 0.89 0.98 1.00
Tg(tba1c:mNeonGreen)cj1-gt 2
0.62 0.69 0.66 0.88 0.88 0.88 0.88 1.00 0.98
Tg(tba1c:mNeonGreen)cj1-gt 1
0.65 0.74 0.69 0.98 0.98 0.98 1.00 0.88 0.89
Tg(actin1:GFP)rs3-in 3
0.65 0.74 0.70 0.98 0.98 1.00 0.98 0.88 0.90
Tg(actin1:GFP)rs3-in 1
0.65 0.74 0.70 0.98 1.00 0.98 0.98 0.88 0.89
Tg(actin1:GFP)rs3-in 4
0.66 0.74 0.71 1.00 0.98 0.98 0.98 0.88 0.90
Tg(actin1:GFP)rs3-in 2
0.92 0.92 1.00 0.71 0.70 0.70 0.69 0.66 0.68
AEP2
0.89 1.00 0.92 0.74 0.74 0.74 0.74 0.69 0.71
AEP1
1.00 0.89 0.92 0.66 0.65 0.65 0.65 0.62 0.63
AEP3
0.0
0.2
0.4
0.6
0.8
1.0

**Figure S14. High reproducibility of neuron-enriched ATAC-seq libraries.** Correlation plot showing the reproducibility of ATAC-seq samples. Spearman correlation analysis demonstrates near-identical profiles across biological replicates. For this study, six neuron-enriched ATAC-seq libraries were generated: two from the *Tg(tba1c:mNeonGreen)*$^{cj1\text{-}gt}$ transgenic line and four from the *Tg(actin1:GFP)*$^{rs3\text{-}in}$ transgenic line (Keramidioti et al., 2024). Additionally, three whole animal ATAC-seq libraries (AEP1-3) from Cazet et al., 2023 were used as controls and benchmarks for data quality standards.

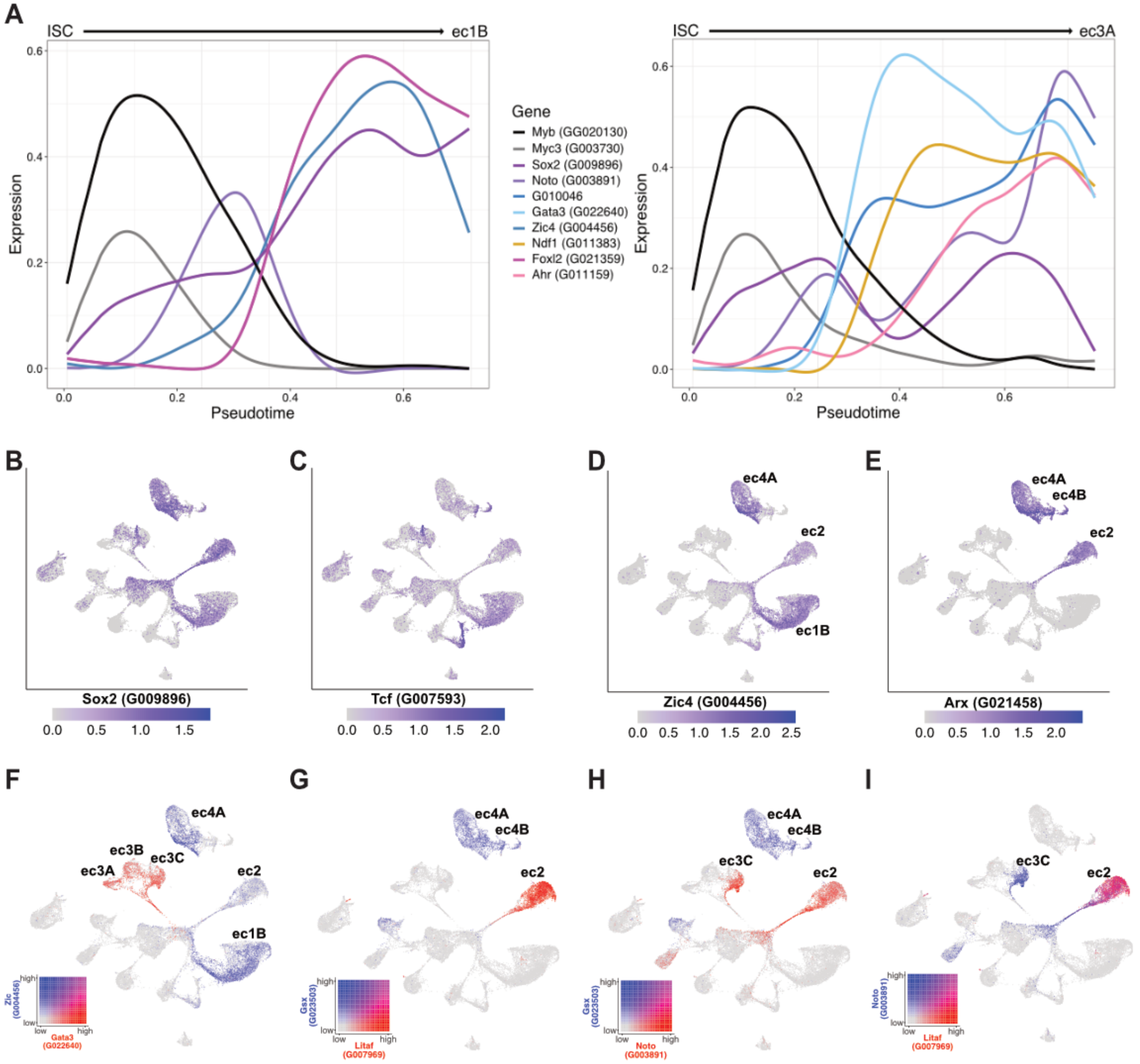
A
ISC
ec1B
Expression
Pseudotime
Gene
Myb (GG020130)
Myc3 (G003730)
Sox2 (G009896)
Noto (G003891)
G010046
Gata3 (G022640)
Zic4 (G004456)
Ndf1 (G011383)
Foxl2 (G021359)
Ahr (G011159)
ISC
ec3A
Expression
Pseudotime
B
Sox2 (G009896)
C
Tcf (G007593)
D
ec4A
ec2
ec1B
Zic4 (G004456)
E
ec4A
ec4B
ec2
Arx (G021458)
F
ec4A
ec3B
ec3C
ec3A
ec2
ec1B
Zic (G004456)
Gata3 (G022640)
G
ec4A
ec4B
ec2
Gsx (G023503)
Litaf (G007969)
H
ec4A
ec4B
ec3C
ec2
Gsx (G023503)
Noto (G003891)
I
ec3C
ec2
Noto (G003891)
Litaf (G007969)

**Figure S15. Transcription factor expression in oral neurons. (A)** Spline plots showing transcription factor expression dynamics during ec1B (left) and ec3C (right) differentiation. **(B–E)** UMAPs showing expression of individual transcription factors: (B) *Sox2* (*G009896*), (C) *Tcf* (*G007593*), (D) *Zic4* (*G004456*), and (E) *Arx* (G021458). **(F-I)** Co-expression UMAPs of the following transcription factor pairs: **(F)** *Gata3* (*G022640*, red) and *Zic4* (blue), **(G)** *Litaf* (G007969, red) and *Gsx1/2* (G023503, blue), **(H)** *Gsx1/2* (red) and *Noto* (*G003891*, blue), **(I)** *Litaf* (red) and *Noto* (blue). Plots in this figure are made using the scRNA-seq portal (https://research.nhgri.nih.gov/HydraAEP/SingleCellBrowser/).

### Supplementary Table Legends

**Table S1. Preparation of neuron-enriched scRNA-seq libraries.** Four neuron-enriched scRNA-seq libraries were generated for this study: one from the *Tg(tba1c:mNeonGreen)*$^{cj1\text{-}gt}$ transgenic line and three from the *Tg(actin1:GFP)*$^{rs3\text{-}in}$ transgenic line (Keramidioti et al., 2024). **Library**: sample type and biological replicate. **Animals Sorted**: Number of animals dissociated and sorted via FACS. **Days of Starvation**: Duration (in days) of starvation prior to dissociation and sorting. **FACS Collection Time (min)**: Time (in minutes) required for FACS to collect the reported number of cells. **GFP+ Cells Collected**: Number of GFP+ cells collected during FACS. **# of Cells Sequenced**: Total number of cells captured and sequenced per library using the Chromium Next GEM Single Cell 3' kit v3.1 (10x Genomics).

**Table S2. Summary metrics for scRNA-seq libraries used in this study.** The study used 18 scRNA-seq libraries: 4 neuron-enriched libraries were generated as part of this study and 14 libraries obtained from Siebert et al., 2019. **Library**: The sample type and biological replicate. **Data From**: Indicates whether the library was generated for this study or obtained from Siebert et al. 2019. **Sequencing Method**: The method used for scRNA-seq collection. **# Cells Sequenced**: The total number of cells captured and sequenced in each library prior to filtering or subsetting. **# Cells Used in Study**: The number of cells included in downstream analyses after filtering and/or subsetting. **Median UMI**: The median number of unique molecular identifiers (transcripts) per cell in each library. **Median Gene**: The median number of unique genes detected per cell in each library.

**Table S3. Median genes and UMIs per cell for each cell state.** This table provides metrics for 19 transcriptionally distinct clusters that were annotated in the study.  **Cell Type**: The cell type annotation based on specific molecular markers. **# Cells**: The total number of cells sequenced for each cell subtype. **Median UMI**: The median number of unique molecular identifiers (transcripts) per cell in each library. **Median Gene**: The median number of unique genes detected per cell in each library.

**Table S4: Neuropeptides found in *Hydra vulgaris*.** This table lists the neuropeptides analyzed in our dot plot (**Fig. S7**), including the name used in our dataset (Column A), the corresponding gene name (Column B), gene identifier (G#; Column C), transcript identifier in the *Hydra vulgaris* AEP transcriptome (t#aep; Column D), and relevant citation(s) (Column E). Gene and transcript identifiers correspond to the annotations in the Cazet et al. (2023) genome assembly and the transcript number correspond to the transcriptome in Siebert et al. (2019).

**Table S5. Details of ATAC-seq library preparation.** This table summarizes the preparation of six neuron-enriched ATAC-seq libraries generated for this study: two from the *Tg(tba1c:mNeonGreen)*$^{cj1\text{-}gt}$ transgenic line and four from the *Tg(actin1:GFP)*$^{rs3\text{-}in}$ transgenic line (Keramidioti et al., 2024). **Library**: The sample type and biological replicate. **Animals Sorted**: Number of animals dissociated and sorted via FACS. **Days of Starvation**: Duration (in days) of starvation prior to dissociation and sorting. **FACS Collection Time (min)**: Time (in minutes) required for FACS to collect the reported number of cells. **GFP+ Cells Collected**: Number of GFP+ cells collected during FACS. **Tagmentation Protocol Time (min)**: Total duration (in minutes) of the ATAC-seq tagmentation protocol (Buenrostro et al., 2015; Corces et al., 2017). **Total PCR Cycles for Amplification**: Number of PCR cycles each library underwent during initial amplification.

**Table S6. Quality control of neuron-enriched ATAC-seq libraries.** This table summarizes the quality control (QC) metrics for six neuron-enriched ATAC-seq libraries generated for this study: two from the *Tg(tba1c:mNeonGreen)*$^{cj1\text{-}gt}$ transgenic line and four from the *Tg(actin1:GFP)*$^{rs3\text{-}in}$ transgenic line (Keramidioti et al., 2024). Additionally, three whole animal ATAC-seq libraries (AEP1-3) from Cazet et al., 2023, were included as controls and benchmarks for data quality. **Library**: The sample type and biological replicate. **Data From**: Indicates whether the library was generated for this study or obtained from Cazet et al., 2023. **Total Read Pairs**: Number of raw read pairs generated for each library. **Final Mapped Read Pairs**: Number of read pairs remaining after the removal of duplicates, unmapped reads, and ambiguously mapped reads. **Alignment Rate**: The percentage of final mapped read pairs that aligned to the reference genome out of the total number of read pairs. According to ENCODE, alignment rates over 80% are considered acceptable (encodeproject.org/atac-seq) (Landt et al., 2012).

**Transcription Start Site (TSS) Enrichment**: Represents the fold enrichment of ATAC-seq signal at the TSS relative to flanking regions (±1 kb). ENCODE sets an acceptable threshold for TSS scores >5, although this varies based on the model system and reflects the biological context. **Reproducible Peaks**: The number of peaks within each library that were reproducibly identified in at least two pairwise comparisons using an irreproducible discovery rate (IDR) cutoff of 0.1. ENCODE considers a threshold of >50,000 reproducible peaks as acceptable. **Self-Consistency Ratio**: A measure of peak consistency within a single dataset. **Rescue Ratio**: The consistency between different datasets. ENCODE considers ideal self-consistency ratios and rescue ratios to be <2.

**Table S7. Primer sequences used for transgenic line construction and fluorescence in situ hybridization (FISH).** This table lists the forward and reverse primers used for amplification of *Hydra* gene regions for either transgenic line generation or FISH probe synthesis. Column A indicates the gene name or annotation used in the study, followed by the forward (Column C) and reverse (Column D) primer sequences. Column E describes the application of each primer pair, including validation of neuronal transition states or spatial localization of specific neuronal subtypes.

**Table S8. Probe pairs used for in situ hybridization chain reaction (HCR) targeting gene *G003886* to label ec1E neurons.**
